# Astrocytes close a critical period of motor circuit plasticity

**DOI:** 10.1101/2020.05.15.098608

**Authors:** Sarah D. Ackerman, Nelson A. Perez-Catalan, Marc R. Freeman, Chris Q. Doe

## Abstract

Critical periods – brief intervals where neural circuits can be modified by sensory input – are necessary for proper neural circuit assembly. Extended critical periods are associated with neurodevelopmental disorders, including schizophrenia and autism; however, the mechanisms that ensure timely critical period closure remain unknown. Here, we define the extent of a critical period in the developing *Drosophila* motor circuit, and identify astrocytes as essential for proper critical period termination. During the critical period, decreased activity produces larger motor dendrites with fewer inhibitory inputs; conversely, increased motor neuron activity produces smaller motor dendrites with fewer excitatory inputs. Importantly, activity has little effect on dendrite morphology after critical period closure. Astrocytes invade the neuropil just prior to critical period closure, and astrocyte ablation prolongs the critical period. Finally, we use a genetic screen to identify astrocyte-motor neuron signaling pathways that close the critical period, including Neuroligin-Neurexin signaling. Reduced signaling destabilizes dendritic microtubules, increases dendrite dynamicity, and impairs locomotor behavior, underscoring the importance of critical period closure. Previous work defines astroglia as regulators of plasticity at individual synapses; here, we show that astrocytes also regulate large-scale structural plasticity to motor dendrite, and thus, circuit architecture to ensure proper locomotor behavior.

## Main

Critical periods are brief windows where neural circuit activity can modify the morphological properties of neurons^1,2^. Critical periods integrate two opposing forces of plasticity to modify neural circuits. <u>Hebbian plasticity</u> alters the function of individual synapses^3^, whereas <u>homeostatic plasticit</u>y encompasses changes to synaptic number, structure (*homeostatic structural plasticity*), and function (*homeostatic synaptic plasticity*) across an entire neuron, as well changes to local and long-range connectivity^1^. While homeostatic plasticity can occur in some areas of the adult brain, dramatic activity-dependent remodeling is largely restricted to early development^3-6^. Indeed, failure to terminate critical period plasticity is linked to a number of neurodevelopmental and neuropsychiatric disorders, such as autism and schizophrenia^2,7-10^. Although critical period closure must be tightly regulated, the molecular mechanisms involved are poorly understood.

### A critical period of motor circuit plasticity

To investigate critical period closure, we focused on two well-characterized *Drosophila* motor neurons (MNs), aCC and RP2^11,12^. These MNs are segmentally repeated in the embryonic and larval CNS (Fig. 1a), and are susceptible to activity-induced remodeling, but these pioneering studies used chronic activity manipulations and did not define an end-point for homeostatic plasticity^12-15^. Here, we expressed the anion channelrhodopsin GtACR2^16^ specifically in the aCC/RP2 MNs and delivered acute 1 hour (h) pulses of silencing terminating at progressively later times in larval development (Fig. 1b-g). We found that silencing MNs in late embryo (stage 17) produced a significant increase in aCC/RP2 dendritic volume at 0 h after larval hatching (ALH). Silencing at later stages showed progressively less of an effect, with no remodeling occurring at 8 h ALH or beyond (Fig. e-g; quantified in 1k). In contrast, acute pulses of activation using the channelrhodopsin Chrimson^17^ resulted in significant loss of MN dendrites at 0 h ALH (Fig. 1h; quantified in Fig. 1k and Extended Data Fig. 1); activating at 8 h ALH and beyond had little or no effect (Fig. 1i-j; quantified in Fig. 1k). Similar results were observed using TrpA1 to thermogenetically activate the MNs (Extended Data Fig. 1). Note that these experiments used far shorter periods of tonic activation than past studies ^13-16,18-21^.

**Figure 1.**
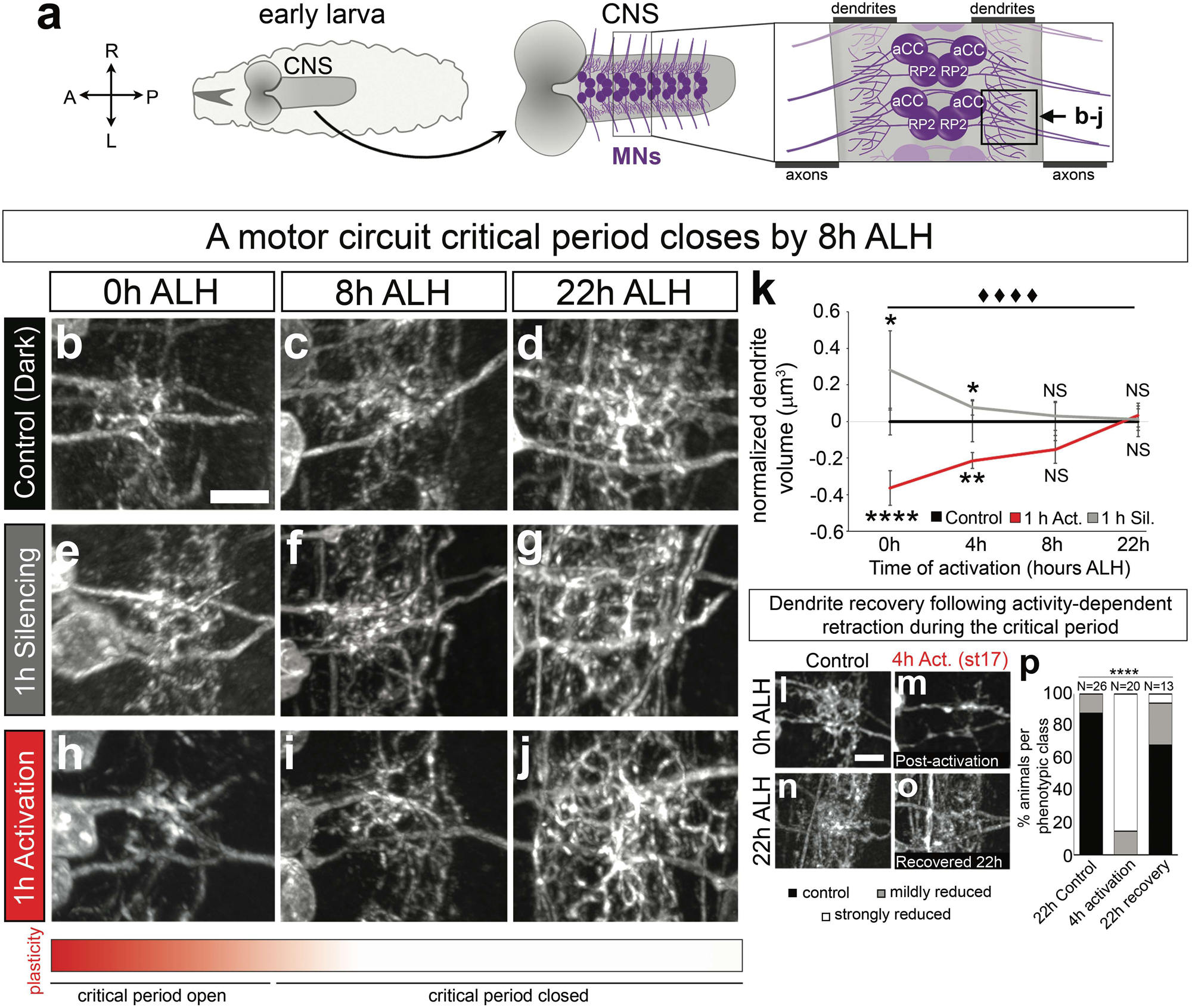
A critical period for motor circuit plasticity at the embryo/larval transition. (**a**) Schematic for reader orientation. A, anterior. P, posterior. L, left. R, right. CNS, central nervous system. MNs, motor neurons. (**b-j**) aCC/RP2 dendritic arbor (single hemisegment) from (**b-d**) dark-reared control, and following (**e-g**) 1 h of light silencing or (**h-j**) 1 h of light activation ending at the indicated stage. Genotypes: Control and silencing: *RN2-gal4,UAS-GtACR2::EYFP*; activation: *RN2-gal4,UAS-CsChrimson::mCherry*. N≥6 brains each, volume averaged across 4 hemisegments (A1-A2). Scale bar, 5 µm. (**k**) Quantification of critical period plasticity. (**l-o**) aCC/RP2 dendritic arbor (single hemisegment) following embryonic activation (st17) and subsequent dark-rearing to allow recovery (0 h *vs*. 22 h ALH). Genotype: *RN2-gal4,UAS-CsChrimson::mCherry*. Brains were categorized qualitatively as in Extended Data Fig. 1**b-d**. (**p**) Quantification. Scale bars, 5 µm. Labels used here and below: *, p<.05; **, p<.01; ***, p<.001; ****, p<.0001, NS= not significant. Error bars: standard deviation and one-way ANOVA used unless otherwise noted. ♦ used in place of * to denote significance following two-way ANOVA when both one-way and two-way are displayed together.

Importantly, activity-induced dendrite loss in late embryo could be rescued by a 22 h period of dark-rearing lacking Chrimson-induced activity (Fig. 1l-p), indicating that activity induces dendrite plasticity, and not excitotoxicity. Together, these experiments define a critical period for activity-dependent motor dendrite plasticity in the early larva, and to our knowledge, represent the first analyses of motor circuit critical period closure within the CNS ^19,22-24^.

### MN activation during the critical period induces dendrite retraction within minutes

In vertebrates, homeostatic plasticity functions on a slow timescale - hours to days^25^. To determine the timescale for MN dendrite expansion following GtACR2 silencing, we silenced aCC/RP2 MNs for three difference lengths of time (15’, 1 h, 4 h) in late embryo and visualized dendritic morphology in single, well-spaced RP2 neurons in newly hatched larvae (0 h ALH) using MCFO^26^. We observed increased dendritic arbor size and complexity following 1 and 4 h of silencing (Fig. 2a-f). We confirmed these results using a different method of neuronal silencing: the dominant negative, temperature sensitive isoform of *shibire*^27^ (Extended Data Fig. 2). In contrast, Chrimson activation resulted in decreased dendrite length and complexity in as little as 15’ activation (Fig. 2g-l). The fact that silencing required more time to show an effect is not altogether surprising, as activity-induced retraction could be achieved through rapid collapse of dynamic cytoskeletal elements, whereas extension requires generation of new membrane^28-31^.

**Figure 2.**
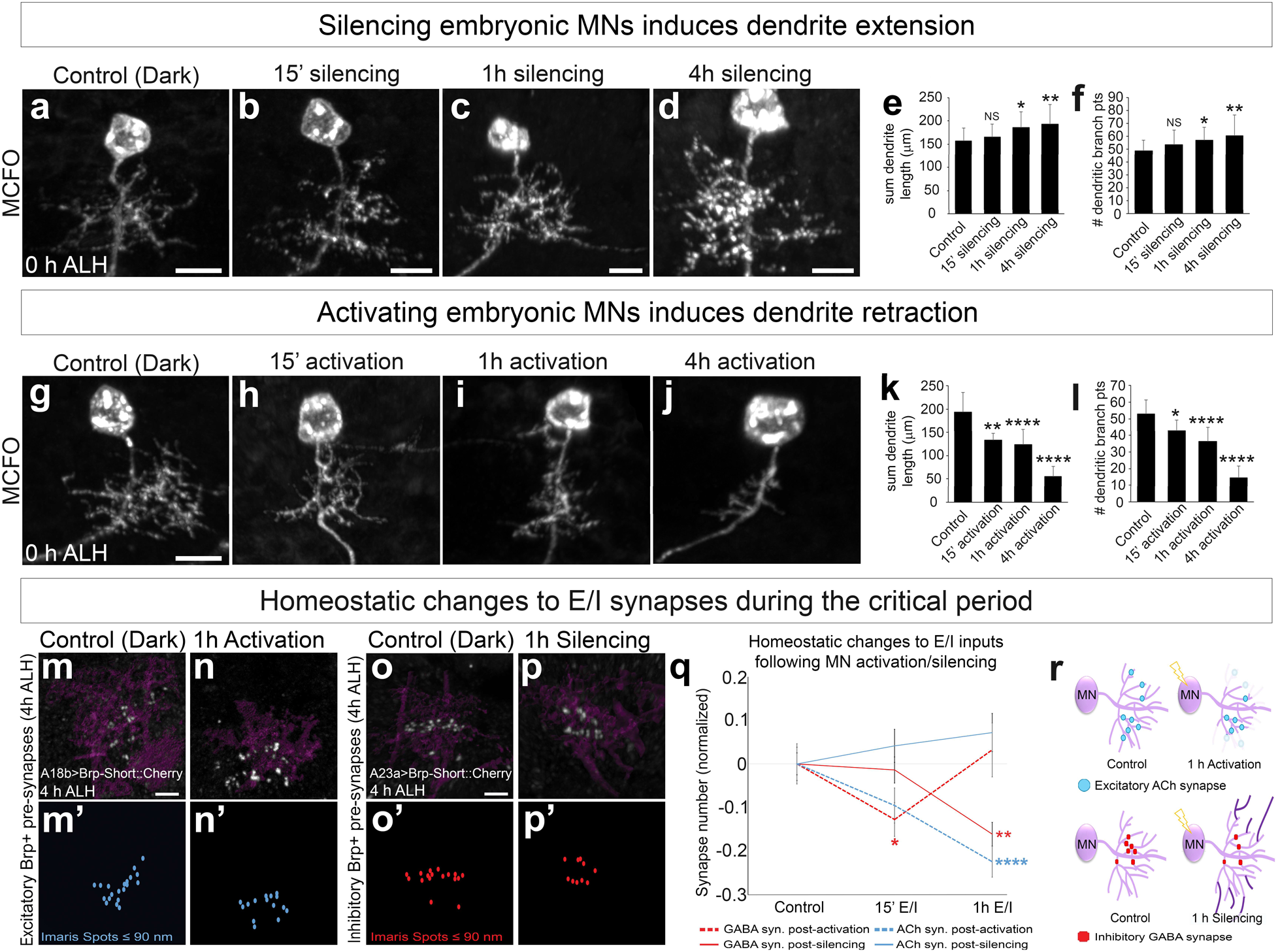
Embryonic MNs scale dendrite length and synaptic inputs to their level of activity. (**a-d**) GtACR2 silencing (or control) for the indicated times prior to 0 h ALH combined with MCFO to visualize single RP2 dendrites; N = #neurons/#animals: N=21/15, 12/9, 13/9, 15/9, respectively. Genotype: *RN2-gal4,UAS-GtACR2::EYFP,UAS-hsMCFO*. Scale bar, 5 μm. (**e-f**) Quantification of dendritic length or branching. (**g-j**) Chrimson activation (or control) for the indicated times prior to 0 h ALH combined with MCFO to visualize single RP2 dendrites; N = #neurons/#animals: 18/15, 7/6, 16/11, 29/19, respectively. Genotype: *RN2-gal4,UAS-CsChrimson::mCherry,UAS-hsMCFO*. Scale bar, 5 μm. (**k-l**) Quantification of dendritic length or branching. (**m-n**) Imaris “Surface” from (**m**) control or (**n**) post-Chrimson activation from 3-4 h ALH (critical period open; magenta, dendrite marker) with presynaptic Brp-short::Cherry puncta (white) from the excitatory A18b neuron; (**m’-n’**) Imaris “Spots”, presynaptic Brp puncta within 90 nm of dendritic surface. Scale bar, 2 µm. Genotype: *RN2-gal4,UAS-Chrimson::mVenus; 94E10-lexA,lexAop-brp-short::cherry*. (**o-p**) Imaris “Surface” from (**o**) control or (**p**) post-GtACR2 silencing from 3-4 h ALH (critical period open; magenta, dendrite marker) with presynaptic Brp-short::Cherry puncta (white) from the inhibitory A23a neuron; (**o’-p’**) Imaris “Spots”, presynaptic Brp puncta within 90 nm of dendritic surface. Scale bar, 2 µm. Genotype: *RN2-gal4,UAS-GtACR2::eYFP; 78F07-lexA,lexAop-brp-short::cherry*. (**q**) Quantification of synapse number following MN excitation or inhibition. N = #hemisegments/#animals: A18b Chrimson N= 18/6 (control); 19/8 (15’ activation); 21/6 (1 h activation). A18b GtACR2 N= 12/9 (control); 17/9 (15’ silencing); 17/8 (1 h silencing). A23a Chrimson N= 33/11 (control); 30/9 (15’ activation); 22/5 (1 h activation). A23a GtACR2 N= 52/13 (control); 36/10 (15’ silencing); 47/17 (1 h silencing). Error bars, SEM. (**r**) Summary.

To further characterize the rapid activation-induced changes in dendrite morphology, we performed live imaging. Both control (myr::GFP) and activated (Chrimson::mVenus) dendrites showed numerous filopodial protrusions over time (Extended Data Fig. 3, Supplementary Movies 1-2), consistent with *in vivo* dendrite dynamics in other systems^32-35^. We first observed significant distal dendrite retraction within 12’ of Chrimson activation (Extended Data Fig. 3d). We conclude that activity-induced remodeling of *Drosophila* MNs occurs on the scale of whole dendritic branches and operates on a time course of minutes, much faster than previously documented for homeostatic plasticity in mammals^25^.

### Activity level scales excitatory/inhibitory synaptic inputs during the critical period

We have shown above that MN silencing increases dendritic arbor size, whereas MN activation decreases arbor size. An important question is whether these morphological changes are accompanied by changes in excitatory or inhibitory (E/I) synaptic inputs. To identify and quantitate E/I inputs, we used the excitatory cholinergic neuron A18b and the inhibitory GABAergic neuron A23a, which we show are each synaptically coupled to the aCC/RP2 dendrites in a TEM reconstruction of the larval CNS^17,36^ (Extended Data Fig. 4). To quantitate E/I synapse number by light microscopy, we used the LexA/LexAop binary system to express a functionally-inactive pre-synaptic marker Bruchpilot^short^::Cherry (Brp)^37^ in A18b or A23a. We quantified Brp puncta overlapping with aCC/RP2 dendritic membrane (putative synapses) using published methods^37-39^ and observed Brp puncta numbers matching synapse numbers by TEM in stage-matched brains (4 h ALH; A18b: 19.5±4.9 Brp+ puncta *vs*. 20±2.5 TEM synapses per hemisegment; A23a: 16.9±4.1 *vs*. 19.5±3.5). Thus, Brp+ puncta contacting MN dendritic membrane are a good proxy for excitatory (A18b) or inhibitory (A23a) premotor synapses.

We quantified A18b excitatory cholinergic synapse number onto aCC/RP2 dendrites before and after activation or silencing. We found that MN activation, but not silencing, significantly decreased A18b excitatory synapses onto aCC/RP2 dendrites (Fig. 2 m-n’; quantified in 2q). Thus, increasing MN activity leads to a compensatory reduction of excitatory pre-synaptic inputs. We next quantified A23a inhibitory GABAergic synapses onto aCC/RP2 dendrites. We found that MN silencing, but not activation, reduced the number of inhibitory synapses between A23a and aCC/RP2 dendrites (Figure 2o-p’; quantified in 2q). Thus, decreasing MN activity leads to a compensatory reduction of inhibitory pre-synaptic inputs. In sum, MNs scale E/I inputs relative to their level of activity during the critical period, presumably to maintain E/I homeostasis (Fig 2r).

### Astrocytes terminate the critical period

Despite the prevalence of well-characterized critical period models in vertebrate systems, the molecular mechanisms that close critical periods are poorly defined. *Drosophila* astrocytes begin to infiltrate the neuropil in the late embryo^40^, prior to closure of the critical period. To test whether astrocytes promote critical period closure, we genetically ablated all astrocytes (see Methods) and used Chrimson to assay for extension of critical period plasticity at 8 h ALH (Fig. 3, Extended Data Fig. 5). Astrocyte elimination was confirmed by staining for the astrocyte marker Gat (Extended Data Fig. 6). As expected, controls closed the critical period by 8 h ALH (Fig. 3a,b; quantified in 3e). In contrast, astrocyte ablation extended the critical period through 8 h ALH (Fig. 3c,d; quantified in 3e). Similar results were observed following 4 h, 1 h, or 15’ of activation (Extended Data Fig. 5). We conclude that astrocytes are required for proper critical period closure. Supporting this conclusion, we found that control motor dendrites were less dynamic after critical period closure, but that astrocyte ablation extends dendrite filopodial dynamicity (Fig. 3g-l, Supplementary Movies 3-6). We conclude that astrocytes are required for the transition from dynamic to stable filopodia, and the concurrent closure of the critical period.

**Figure 3.**
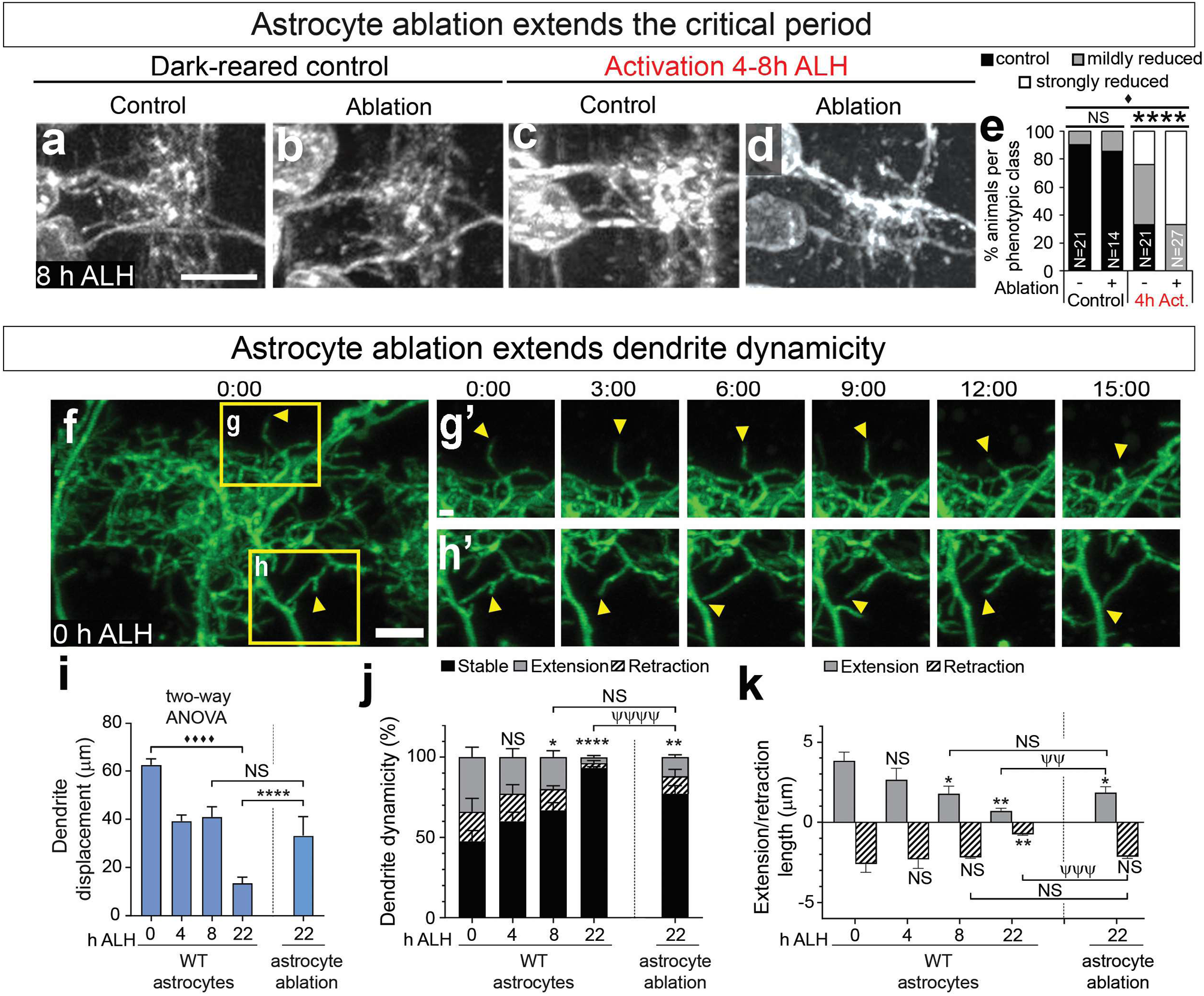
Astrocytes terminate the critical period. (**a-e**) Astrocyte ablation prolongs the critical period. **(a-d)** aCC/RP2 dendrites in two hemisegments at a 8 h ALH. (**a-b**) Dark-reared controls with or without astrocyte ablation. (**c-d**) Chrimson activation in aCC/RP2 from 4-8 h ALH; note that astrocyte ablation prolongs the critical period to allow activity-dependent dendrite reduction. Genotypes: *RN2-gal4,UAS-CsChrimson::mCherry*; *alrm-lexA,lexAop-myr::GFP* (control), *RN2-gal4,UAS-CsChrimson::mCherry*; *alrm-lexA,lexAop-rpr* (ablation). Scale bar, 5 µm. (**e**) Quantification. **(f-k)** Astrocyte ablation prolongs dendrite dynamicity. (**f**) Live imaging of dendrite dynamics. 3D projection, one hemisegment of aCC/RP2 dendrites at 0 h ALH. Yellow boxes (**g-h**), regions followed over time. Scale bar: 5 µm. (**g’-h’**) Dynamic dendrite filopodia (arrowheads) imaged for 15’. Scale bar, 1 µm. Genotypes: *RN2-gal4,UAS-myr::GFP; alrm-lexA* (control), *RN2-gal4,UAS-myr::GFP; alrm-lexA,lexAop-rpr* (ablation). (**i-k**) Quantification. N=50 dendrites from 5 brains per timepoint. ψ: comparisons between ablation and controls (Fisher’s exact tests).

### Identification of astrocyte signaling pathways that close the critical period

How do astrocytes close the critical period? Astrocytes are known to communicate with neurons for proper synapse formation, elimination, and function via both cell surface molecules and secreted proteins^41-43^. We therefore used the astrocyte-specific *alrm-gal4* to perform an RNAi knock down (KD) screen using commercially available *UAS-RNAi* lines^44^. We tested 61 lines encompassing 49 genes curated for known functions in astrocytes and/or genes identified in a parallel screen for astrocyte-derived genes that regulate motor function (see Methods). Animals were reared at 30°C to obtain maximum RNAi expression and assayed at 8 h ALH for extension of the critical period. We assayed Chrimson-induced plasticity, as dendrite retraction is more rapidly screenable by eye. Knockdown of most genes had no effect on critical period closure. However, four genes were required in astrocytes for timely critical period closure: *gat* (regulates E/I balance), *CG43313* (synthesizes inhibitory extracellular matrix CSPGs), and the Neuroligins (Nlg) 4 and 2 (Fig. 4a-g). Importantly, KD of each gene had little or no effect on astrocyte survival or morphology (Extended Data Fig. 6), suggesting a more specific defect in astrocyte-motor neuron signaling.

Here, we focus on Neuroligins, which bind cell adhesion proteins called Neurexins (Nrx). We used RNAi against *nrx-1*, known to bind both *nlg2/4*^45,46^, specifically in aCC/RP2 MNs, and observed extension of the critical period (Fig. 4h-k); this is consistent with astrocyte Nlg2/4 and MN Nrx-1 acting in a common pathway to close the critical period. Notably, while Nrx-1 is generally considered pre-synaptic, there is ample evidence in both invertebrate and vertebrate systems for dendritic localization of these receptors^47-50^. We next used previously published Crispr-induced overexpression lines^51^ for Nrx-1 and Nlg2 to determine if they could induce precocious critical period closure. As expected, controls with Chrimson activation in aCC/RP2 from 3-4 h ALH showed strong dendritic reduction (Fig. 1k); in contrast, forced expression of Nrx-1 in aCC/RP2 MNs prematurely closed the critical period, as seen by absence of Chrimson-induced dendritic loss at 3-4 h ALH (Fig. 4l-m, quantified in 4o). Similarly, overexpression of Nlg2 alone in astrocytes was sufficient to prematurely close the critical period (Fig. 4n-o). We conclude that the Nlg2/Nrx-1 ligand/receptor pair are required in astrocytes and MNs (respectively) for timely closure of the critical period.

**Figure 4.**
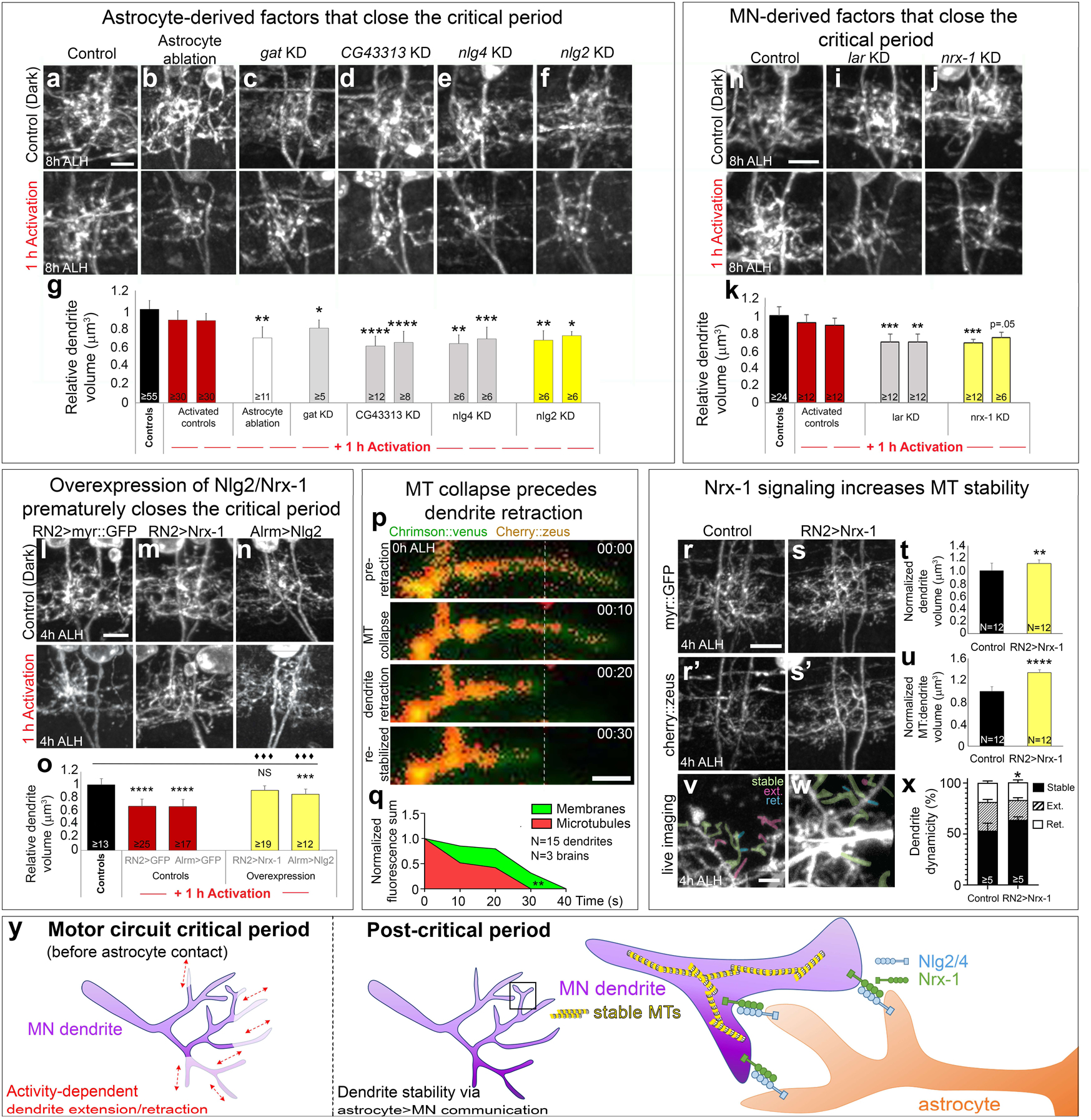
Astrocyte Neuroligin signals to MN Neurexin to stabilize microtubules and close the critical period. (**a-k**) Factors in astrocytes (**a-f**) or MNs (**h-j**) required to close the critical period. aCC/RP2 dendrites in one hemisegment at 8 h ALH. Top row, dark reared controls; bottom row, experimentals. Genotypes: (**a**,**c-f**) *lexAop-CsChrimson::mVenus,RN2-lexA,alrm-gal4 UAS-RNAi*, (**b**) *RN2-gal4,UAS-Chrimson::mCherry,lexAop-rpr, alrm-lexA* (**h-j**), *RN2-gal4,UAS-CsChrimson::mCherry UAS-RNAi*. Scale bar, 5 µm. (**g**,**k**) Quantification, two-way ANOVA. (**l-n**) Precocious critical period closure at 4 h ALH by overexpression of (**m**) Nrx-1 or (**n**) Nlg2 in MNs or astrocytes, respectively. Genotypes: (**l-m**) *RN2-gal4,UAS-CsChrimson::mCherry* x *UAS-myr::GFP* or *UAS-Nrx-1*, (**n**) *RN2-lexA,lexAop-Chrimson::tdTomato,alrm-gal4 x UAS-Nlg2*. Scale bar, 5 µm. (**o**) Quantification. (**p**) Live imaging of aCC/RP2 dendrites expressing Chrimson::mVenus (green) and Cherry::zeus (stable microtubules, heatmap) at 0 h ALH. Dashed line, retraction landmark. (**q**) Quantification, Two-way ANOVA. (**r-s**) Dendritic (myr::GFP) distribution of (**r’-s’)** microtubules (Cherry::Zeus) in (**r-r’**) controls and (**s-s’**) post-overexpression of Nrx-1 in MNs at 4 h ALH. Genotypes: (**r-r’**) *RN2-gal4,UAS-myr::GFP,UAS-Cherry::Zeus,UAS-redstingerNLS* (**s-s’**) *RN2-gal4,UAS-myr::GFP, UAS-Cherry::Zeus,UAS-Nrx-1*. (**t-u**) Quantification of dendrite volume or microtubule:dendrite volume. (**v-w**) Live imaging of stable microtubules (Cherry::Zeus+) in aCC/RP2 (**v**) control or (**w**) Nrx-1 overexpression dendrites. Genotypes: (**v**) *RN2-gal4,UAS-myr::GFP,UAS-Cherry::Zeus*, (**w**) *RN2-gal4, UAS-Cherry::Zeus,UAS-Nrx-1*. Pseudocoloring: stable (green), extending (pink), or retracting (blue) dendrites. (**x**) Quantification, Fisher’s Exact Test. (**y**) Summary.

### Nrx-1 signaling stabilizes dendritic microtubules at critical period closure

How does Nlg2/Nrx-1 signaling close the critical period? Nrx-1 promotes motor axon microtubule stability^52,53^, suggesting a microtubule-stabilization mechanism for critical period closure. To test this hypothesis, we used Chrimson::mVenus to activate and visualize aCC/RP2 dendrite membranes at 0 h ALH (peak critical period), and Cherry::Zeus to visualize stable microtubules during and after dendritic retraction. In live preparations, Cherry::Zeus was most robust in proximal dendritic arbors, though stable microtubules were also observed in extending distal processes (Fig. 4p). Interestingly, processes that undergo remodeling showed a reduction in Cherry::Zeus intensity immediately preceding dendrite retraction (Fig. 4q, Supplementary Movie 7), suggesting that microtubule collapse within distal branches can induce dendrite retraction. In fixed preparations, we found that proximal dendrites with the highest levels of stable microtubules were protected from activity-dependent retraction (Extended Data Fig. 7). Interestingly, overexpression of Nrx-1 was sufficient to increase both stable microtubules and dendrite stability (Fig. 4r-x, Supplementary Movies 8-9). We propose that Nlg2 in astrocytes binds Nrx-1 in MNs to stabilize dendritic microtubules and close the critical period (Fig. 4y; see Discussion).

### Timely closure of the critical period is required for normal locomotor behavior

In mammals, inappropriate extension of critical periods compromises nervous system function^2^. We extended the critical period by temperature controlled RNAi of critical period regulators until 12 h ALH, and then restored gene expression until 44 h ALH, when they were assayed for locomotor behavior (protocol established in Extended Data Fig. 8 and illustrated in Extended Data Fig. 9a-b). Control larvae showed strong linear persistence; in contrast, most larvae with extended critical periods showed excessive turning resulting in spiraled trajectories. We also observed deviations in speed, distance from origin, accumulated distance, cumulative bending angle, or pausing in larvae with extended critical periods (Extended Data Fig. 9c-t). We propose that timely closure of the MN critical period is essential for normal larval locomotor behavior.

## Discussion

Astrocytes have a well-characterized role in synaptogenesis, synaptic pruning and synaptic efficacy^51^, but little is known about their role in critical period closure. In this study, we identified astrocytes as promoting closure of a motor critical period required for locomotor function, and define a series of astrocyte-MN signaling pathways, both known and novel, required to close the critical period. Based on previous literature, we hypothesized that astrocytes could modify critical period closure through regulation of E/I balance^2,52^ or extracellular matrix composition^42^. Consistent with mammalian studies, we found that perturbing E/I balance through astrocyte-specific RNAi of the sole GABA transporter, Gat, was sufficient to extend critical period plasticity. Further, we found that increasing levels of extracellular matrix chondroitin sulfate proteoglycans (CSPGs) through RNAi KD *CG43313*, homologous to mammalian Chondroitin sulfate synthase 2 enzyme^53^, extended critical period plasticity. Similarly, MN-specific RNAi KD of the CSPG receptor *lar*^54^ also extended critical period plasticity (Fig. 4h-k). Thus, our data suggest that astrocytes employ similar strategies in both *Drosophila* and mammals to regulate critical periods. Unexpectedly, we also identified astrocyte-derived Neuroligins, and their neuronal partner Nrx-1, as instrumental for critical period closure. In mammals, Neuroligins are known to regulate synapse formation and astrocyte morphology^43^, but their role in regulating critical period closure is novel.

Our data support the hypothesis that Nrx-1 signaling in motor dendrites increases local microtubule stability to close the critical period, but how Nrx-1 alters microtubule stability remains to be tested. Recent reports indicate that local reactive oxygen species (ROS) signaling can trigger homeostatic dendritic retraction^21,55^. Mutations in Neuroligins are associated with increased ROS sensitivity^56^. Further, microtubule-binding proteins are known targets of ROS and increased ROS levels can destabilize microtubules^57^. It is interesting to speculate that during the critical period, rapid dendritic retraction is achieved through local accumulation of ROS, which is suppressed upon Neuroligin-Neurexin signaling from astrocytes to MNs. In sum, closure of the motor circuit critical period is induced by astrocyte Neuroligin to MN Neurexin signaling to stabilize dendritic microtubules.

## Supporting information

Full Supplementary Data, Methods, and Movies

Ackerman2020_SuppMovies

Ackerman2020_SuppData_Methods

## DATA AND CODE AVAILABILITY

This study did not generate/analyze datasets/code.

## Acknowledgements

We thank Takashi Suzuki, Stephen Cohen, Ellie Heckscher, and Vivek Jayaraman for providing fly stocks. We thank Kelly Monk, Jim Skeath, Dave Lyons, and members of the Doe lab for comments on the manuscript. Stocks obtained from the Bloomington Drosophila Stock Center and Shigen National Institute of Genetics (NIH P40OD018537) were used in this study. Funding was provided by HHMI (CQD), R01 HD27056 (CQD), R01 NS059991 (MRF) and NIH F32NS098690 (SDA). SDA is a Milton Safenowitz Post-doctoral fellow of the ALSA.

## Author Contributions

SDA conceived of the project; SDA and NPC performed experiments; MRF and CQD provided feedback during the project; SDA, NPC, and CQD wrote the paper and prepared the Figures. All authors commented and approved of the manuscript.

