## Supplementary material for "Astrocytes close a critical period of motor circuit plasticity": Full Supplementary Data, Methods, and Movies: Ackerman_SupFigures_Methods_15May2020.pdf

### SUPPLEMENTARY MATERIALS: DATA

#### Activity-dependent dendrite retraction following Chrimson-activation

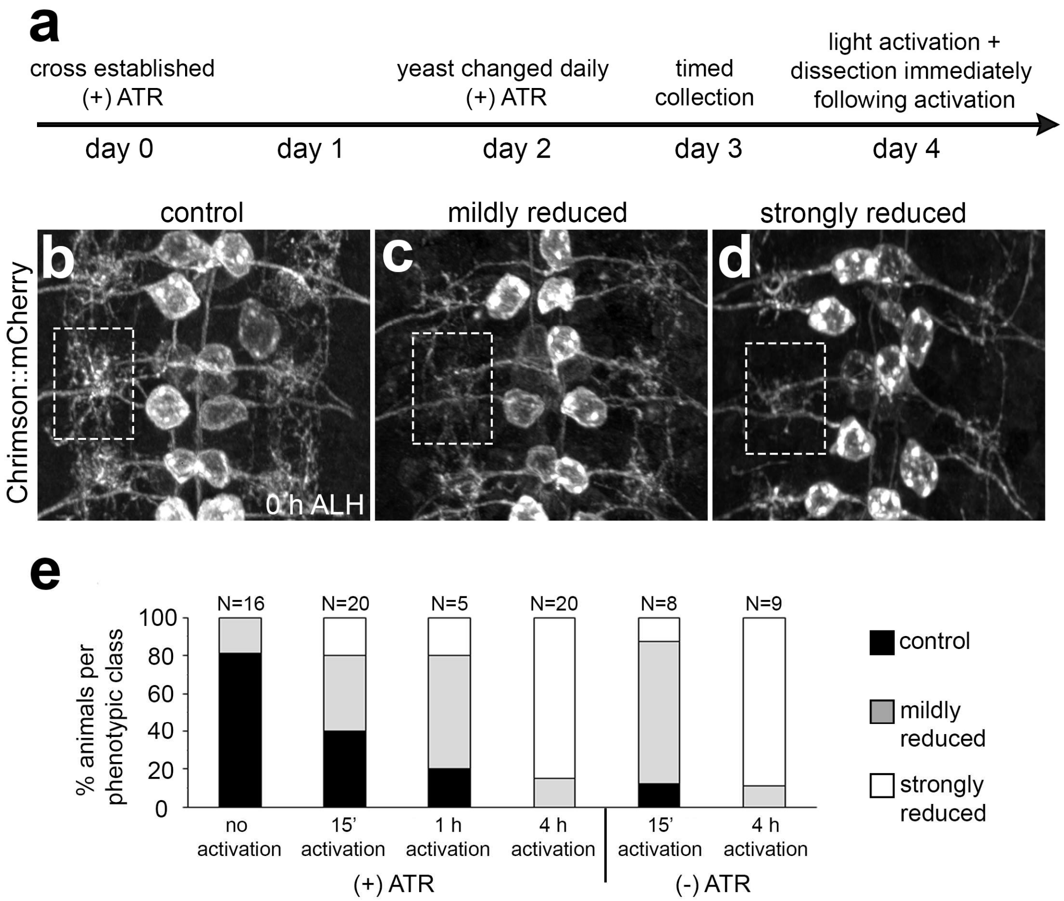


#### Dendrite retraction by TrpA1 activation mimics Chrimson

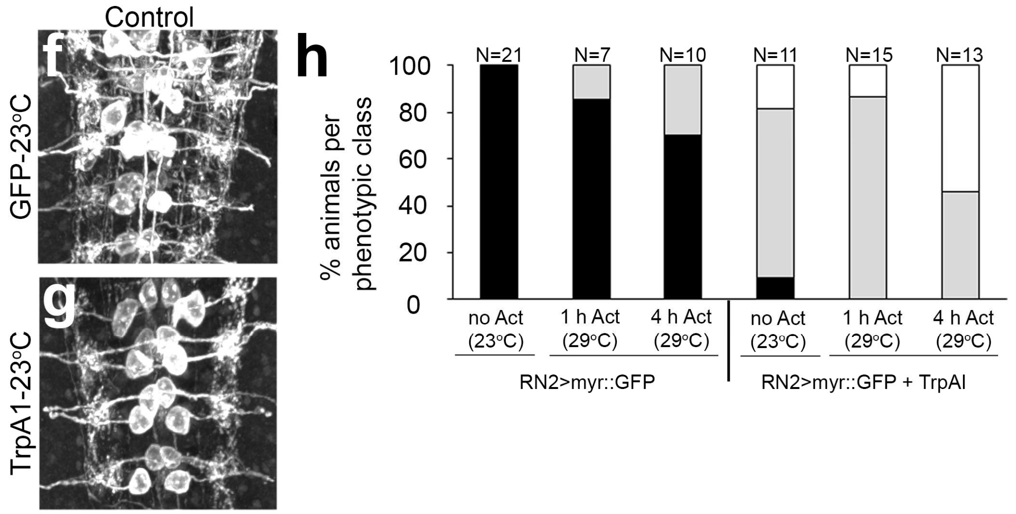

**Extended Data Figure 1. Activation of motor neurons results in rapid retraction of dendrites.**

(**a**) Schematic of the activation paradigm used in this study. For activation of aCC/RP2 motor neurons (MNs), *RN2-gal4/lexA* drove expression of *UAS-CsChrimson::mCherry* or *lexAop/UAS-CsChrimson::mVenus*. Crosses were established on day 0 and fed exclusively on yeast paste supplemented with 0.5 mM all-trans retinal (ATR; required for full Chrimson activity) and changed daily for a minimum of 3 days. Timed embryo collections were performed on day 3 for a duration of 1.5 h. Sustained light activation (10550 lx) was followed by immediate dissection. Optogenetic silencing experiments using *UAS-GtACR2::EYFP* followed the same scheme.
(**b-h**) Activation of aCC/RP2 MNs by Chrimson channelrhodopsin or TrpAI induces dendrite retraction.
(**b-d**) Representative 3D projections of brains expressing Chrimson::mCherry in aCC/RP2 MNs at 0 h after larval hatching (ALH) following activation during embryonic stage 17 (st17). After activation, brains were categorized qualitatively as (**b**) control, (**c**) mildly reduced or (**d**) strongly reduced based on the extent of aCC/RP2 dendritic elaboration (dashed white boxes).
(**e**) Quantification of each phenotypic class in control, dark-reared animals versus animals whose aCC/RP2 MNs were Chrimson-activated for 15’, 1 h, or 4 h. Dark-reared controls were used throughout as aCC/RP2 MNs show sensitivity to Chrimson in the absence of ATR ((-) ATR) after 15’ and 4 h of Chrimson activation.
(**f-h**) Thermogenetic activation of aCC/RP2 (*RN2-gal4 UAS-TrpA1* or *UAS-myr:GFP* control) mimics Chrimson-activation. TrpA1 causes maximum activation at 29°C (≥30 Hz) and is inactive at 23°C. (**g**) Representative TrpA1 control (reared at 23°C) shows reduced dendrite elaboration relative to (**f**) GFP control, suggesting that TrpA1 is partially active at 23°C. (**h**) Quantification of each phenotypic class (**b-d**) in GFP controls reared at 23°C *vs.* 29°C for 1 h or for 4 h, relative to TrpA1 animals reared at 23°C (control), or activated for 1 h or 4 h at 29°C.

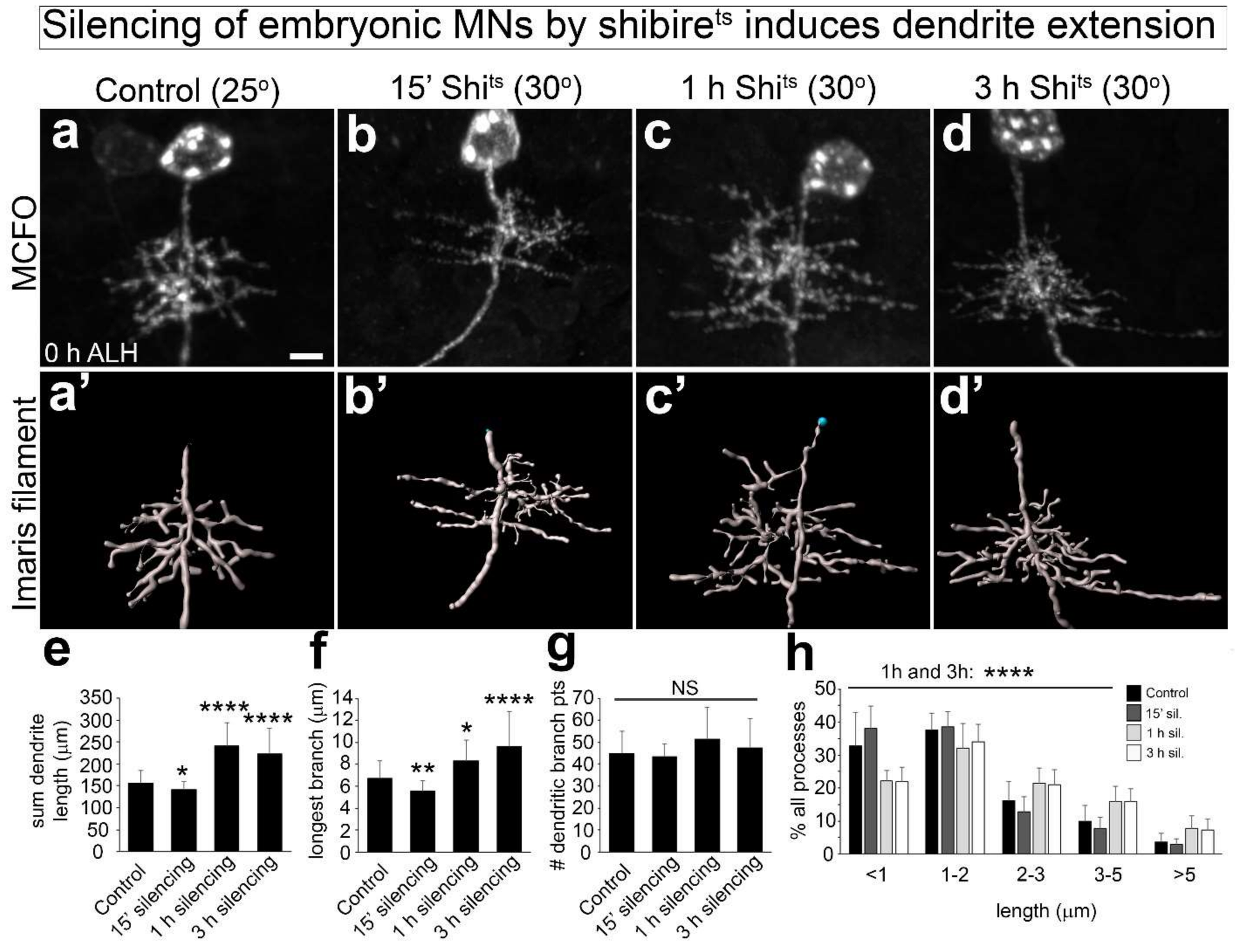


**Extended Data Figure 2. Silencing of RP2 by shibire induces dendrite extension.**
(**a-d**) MCFO single neuron labeling at 0 h ALH to visualize the morphology of RP2 MN dendrites at 25°C in (**a**) shibire control (N=43 neurons/N=24 brains) compared to neurons silenced with temperature sensitive shibire to block synaptic transmission (active at 30° C) for (**b**) 15 minutes (N=18/N=15), (**c**) 1 h (N=7/N=6), or (**d**) 3 h (N=29/N=18). Scale bar, 5 μm. *Prime* panels show reconstructions of RP2 dendritic arbors (performed using the Imaris "Filaments" tool). Blue dots show the seed positions for each Filament.
(**e-h**) Quantification of (**e**) total dendrite length, (**f**) longest branch length (measure of distal dendrite extension), (**g**) # of dendritic branch points, and (**h**) the distribution of dendrite lengths per reconstructed neuron (% of all processes) post-silencing by shibire. (**e**) Total dendrite length, (**f**) longest branch length, and (**h**) the percentage of long processes (>2 μm) were significantly increased after 1 h and 3 h of silencing (subtle decreases after 15'). (**g**) Silencing led to no changes in dendritic complexity, measured by the number of dendritic branch points within a single arbor. From here and below: all statistics represent one-way ANOVA; error bars represent standard deviation; and *, p<.05; **, p<.01; ***, p<.001; ****, p<.0001.

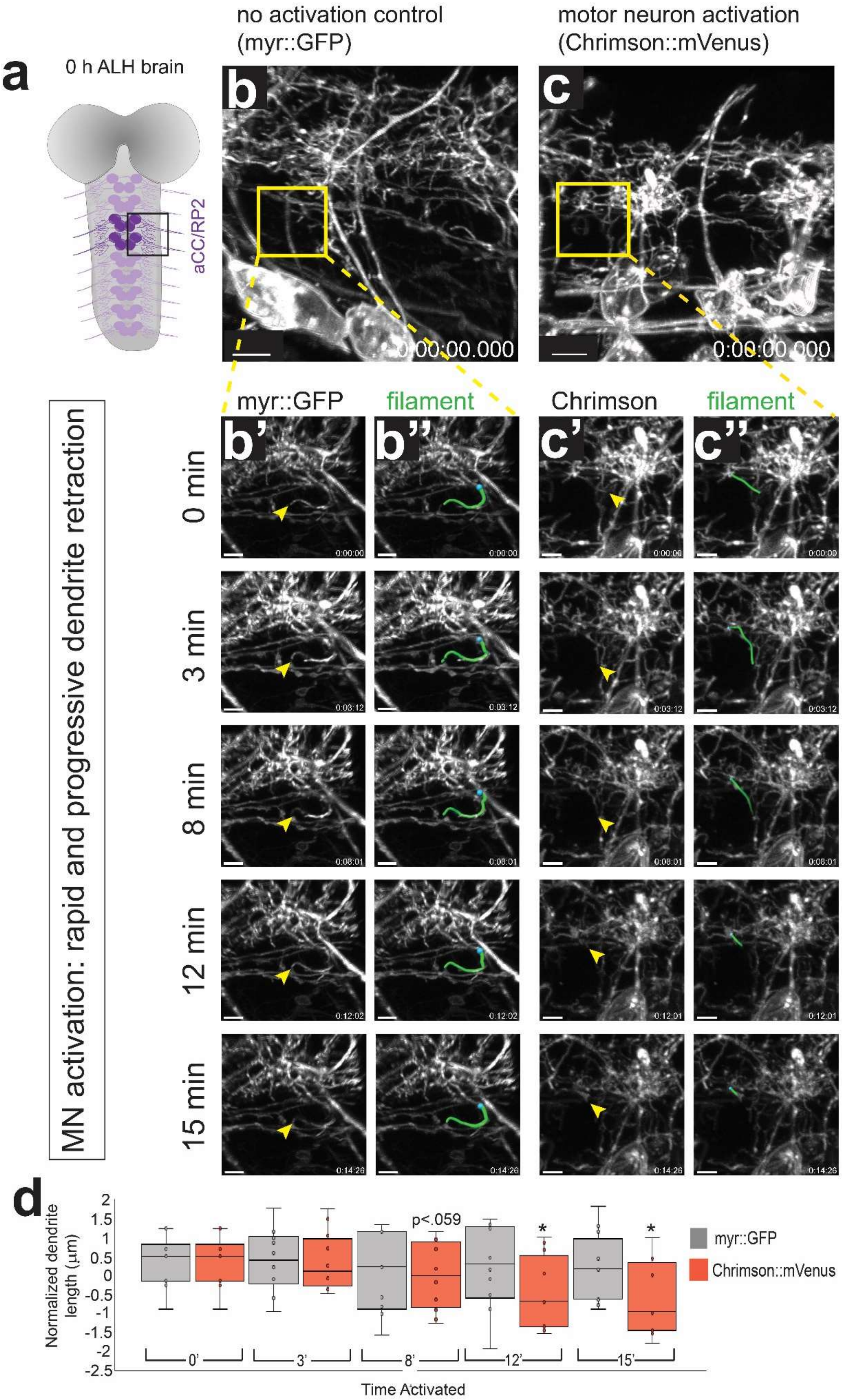

a
0 h ALH brain
aCC/RP2
no activation control
(myr::GFP)
motor neuron activation
(Chrimson::mVenus)
b
c
0:00:00.000
0:00:00.000
MN activation: rapid and progressive dendrite retraction
myr::GFP
filament
Chrimson
filament
b'
b''
c'
c''
0 min
3 min
8 min
12 min
15 min
d
Normalized dendrite length (μm)
p<.059
myr::GFP
Chrimson::mVenus
0'
3'
8'
12'
15'
Time Activated

**Extended Data Figure 3. Remodeling of dynamic distal processes within 12 minutes of aCC/RP2 activation.**
(**a**) Schematic depicting a larval brain at 0 h ALH with aCC/RP2 MNs in purple. Two hemisegments were imaged per experiment (box).
(**b-d**) MN Chrimson activation results in dendrite retraction within minutes.
(**b**) 3D projection of a control isolated CNS at 0 h ALH, time 0 (*RN2-gal4*,*UAS-myr::GFP*; + ATR). Yellow box highlights intersegmental region used for reconstruction of individual dendrites. Scale bar, 5 µm.
(**b'-b"**) 3D projections from representative time points over a 15-minute acquisition period. Left panels, myr::GFP signal alone. Yellow arrowheads mark the tip of a single reconstructed process. Right panels, green Imaris "Filament" reconstruction of indicated process. Scale bars, 1 µm.
(**c**) 3D projection of an isolated CNS at 0 h ALH for Chrimson-activation, time 0 (*RN2-gal4,UAS-CsChrimson::mVenus*; + ATR). Yellow box highlights intersegmental region used for reconstruction of individual dendrites. Scale bar, 4 µm.
(**c'-c"**) 3D projections from representative time points over a 15-minute acquisition period. Left panels, Chrimson::mVenus signal alone. Yellow arrowheads mark the tip of a single reconstructed process. Right panels, green Imaris "Filament" reconstruction of indicated process. Scale bars, 1 µm.
(**d**) Quantification of normalized dendrite length over time in myr::GFP controls versus Chrimson-activated brains (N=10 processes each from N=4 brains per condition, with processes binned by length into 10 categories). Control length remained stable over the 15-minute acquisition period. Chrimson-activation results in progressive retraction of MN dendrites.

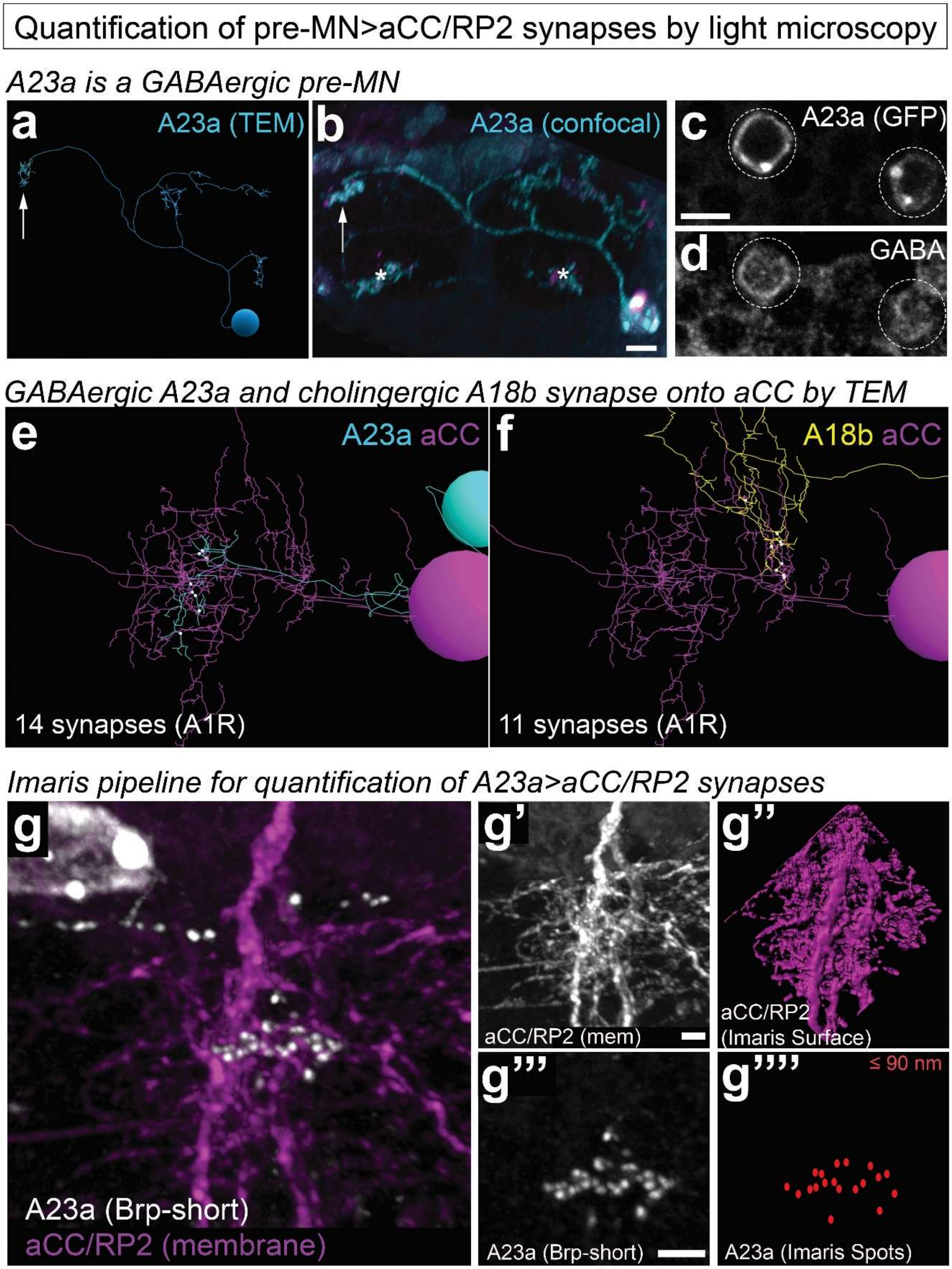

Quantification of pre-MN>aCC/RP2 synapses by light microscopy
A23a is a GABAergic pre-MN
a
A23a (TEM)
b
A23a (confocal)
c
A23a (GFP)
d
GABA
GABAergic A23a and cholingergic A18b synapse onto aCC by TEM
e
A23a aCC
14 synapses (A1R)
f
A18b aCC
11 synapses (A1R)
Imaris pipeline for quantification of A23a>aCC/RP2 synapses
g
A23a (Brp-short)
aCC/RP2 (membrane)
g'
aCC/RP2 (mem)
g''
aCC/RP2
(Imaris Surface)
g'''
A23a (Brp-short)
g''''
≤ 90 nm
A23a (Imaris Spots)

**Extended Data Figure 4. Quantification of the number of synaptic connections between the GABAergic A23a or cholinergic A18b interneurons and the motor neurons aCC/RP2.**
(**a**) TEM reconstruction of the A23a premotor neuron in a first instar larval brain at 4 h ALH; pre-synapses are primarily localized to the contralateral branch (arrow).
(**b**) Light microscopy image of a single A23a premotor neuron at 4 h ALH (*78F07-lexA*) with cyan membranes (*lexAop-myr::GFP*) and magenta pre-synapses (*lexAop-brp-short::cherry*). Most synapses with aCC/RN2 MNs are at the contralateral process (arrow). Note the morphological similarity between light and EM images of A23a. Asterisks, sparse off-target expression not in A23a. Scale bar, 2 µm.
(**c-d**) A23a is GABAergic. Genotype: *78F07-lexA lexAop-myr::GFP*. Scale bar, 3 µm.
(**e**) Representative image of A23a (A1L) forming 14 synapses (white dots) with aCC (A1R) in the TEM reconstruction. Dorsal view, midline to left. Quantification of A23a-aCC synapses from the TEM reconstruction: 21 in A1R and 13 in A1L; A23a-RP2 synapses: 2 in A1R and 3 in A1L.
(**f**) Representative image of A18b (A1L) forming 11 synapses (white dots) with aCC (A1R) in the TEM reconstruction. Dorsal view, midline to left. Quantification of A18b-aCC synapses from the TEM reconstruction: 18 in A1R and 10 in A1L; A18b-RP2 synapses: 7 in A1R and 5 in A1L.
(**g-g""**) Quantification of "putative" A23a-aCC/RP2 synapses by light microscopy at 4 h ALH.
(**g**) Representative 3D projection of aCC/RP2 dendrite membrane (Chrimson::mVenus+; magenta) and A23a Brp-short puncta (white). Genotype: *RN2-gal4,UAS-Chrimson::mVenus* x *78F07-lexA,lexAop-brp-short::cherry*. (**g'**) aCC/RP2 dendrite membrane (Chrimson::mVenus+); (**g"**) Imaris "Surface" rendering of **g'**; (**g'"**) A23a pre-synapses (Brp-short::cherry+); (**g""**) Imaris "Spots" measurement of Brp-short puncta within 90 nm of dendritic membrane (red dots, 19 putative direct synapses).

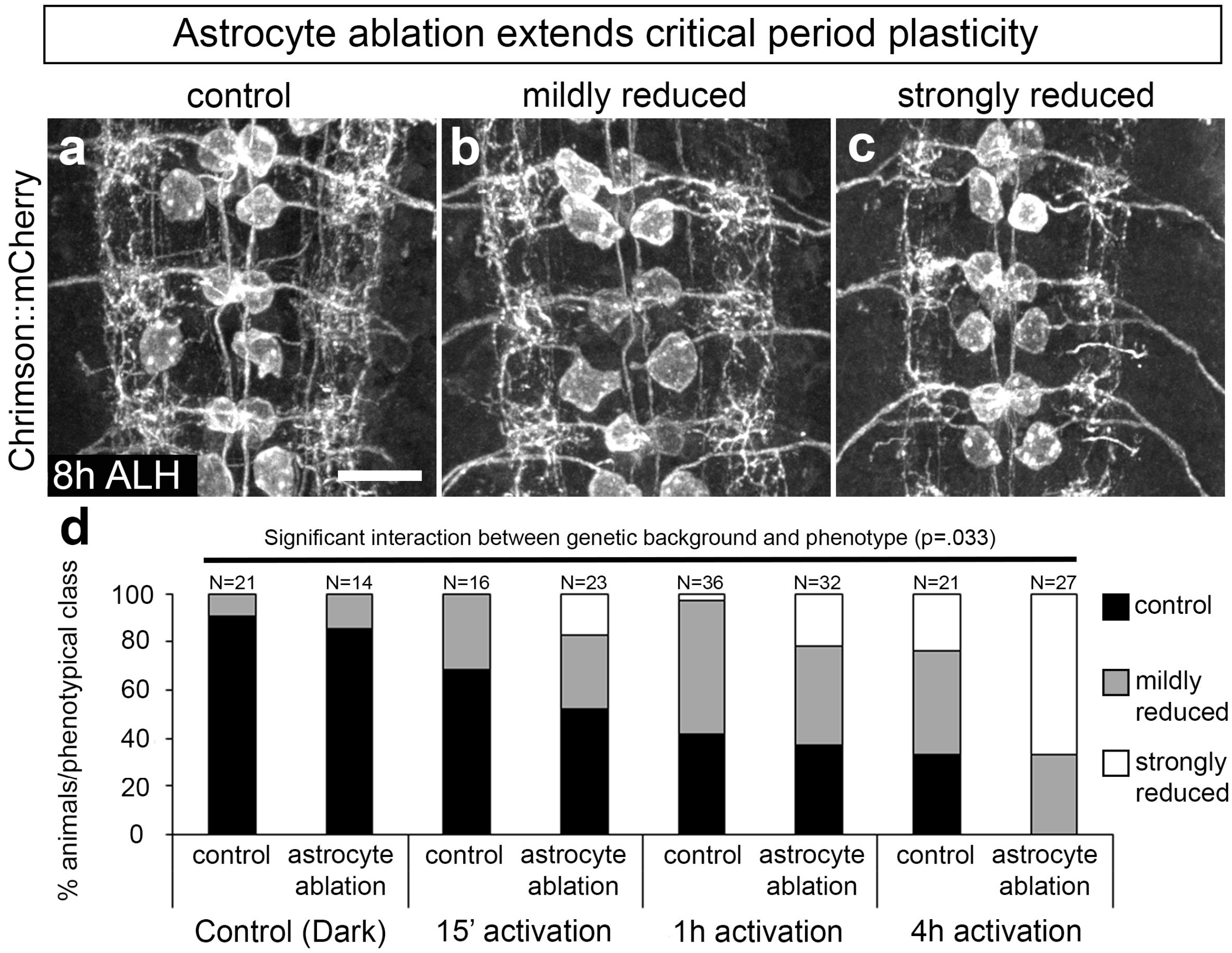


**Extended Data Figure 5. Astrocyte ablation extends critical period plasticity.**
(**a-c**) Representative 3D projections of brains expressing Chrimson::mCherry in aCC/RP2 motor neurons (*RN2-gal4,UAS-Chrimson::mCherry*) illustrating the three classes of dendritic arbor morphology at 8 h ALH following 4 h of Chrimson activation: (**a**) control, (**b**) mildly reduced, and (**c**) strongly reduced dendritic arbor size/complexity. Scale bar, 10 μm.
(**d**) Quantification of each phenotypic class. Control animals show no significant dendritic remodeling after 15' of activation at this stage ($p<.12$, one-way ANOVA). In contrast, ablation (abl.) of astrocytes results in a significant shift in the distribution of phenotypic classes away from wildtype (no light abl. versus 15' activation abl., $p<.03$, one-way ANOVA). Loss of astrocytes strongly sensitized these motor neurons to remodeling ($p<.04$, two-way ANOVA). Note that control and 4 h data are also displayed in Fig. 3e. Control Genotype: *RN2-gal4,UAS-Chrimson::mCherry; alrm-lexA,lexAop-myr::GFP*. Ablation Genotype: *RN2-gal4,UAS-Chrimson::mCherry; alrm-lexA,lexAop-rpr*.

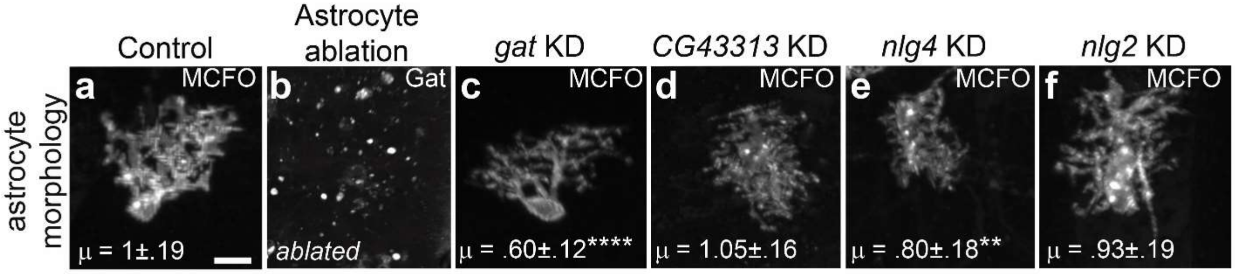


**Extended Data Figure 6. Astrocyte morphology post-knockdown of critical period regulators.**

(**a-f**) MCFO showing single astrocyte morphology and volume, or the pan-astrocyte marker Gat at 8 h ALH. Normalized, mean astrocyte volume at the bottom of each MCFO panel (via Imaris "Surface"), with significance indicated by asterisks (N≥ 12 astrocytes from N≥4 brains per genotype). Scale bars, 5 μm.

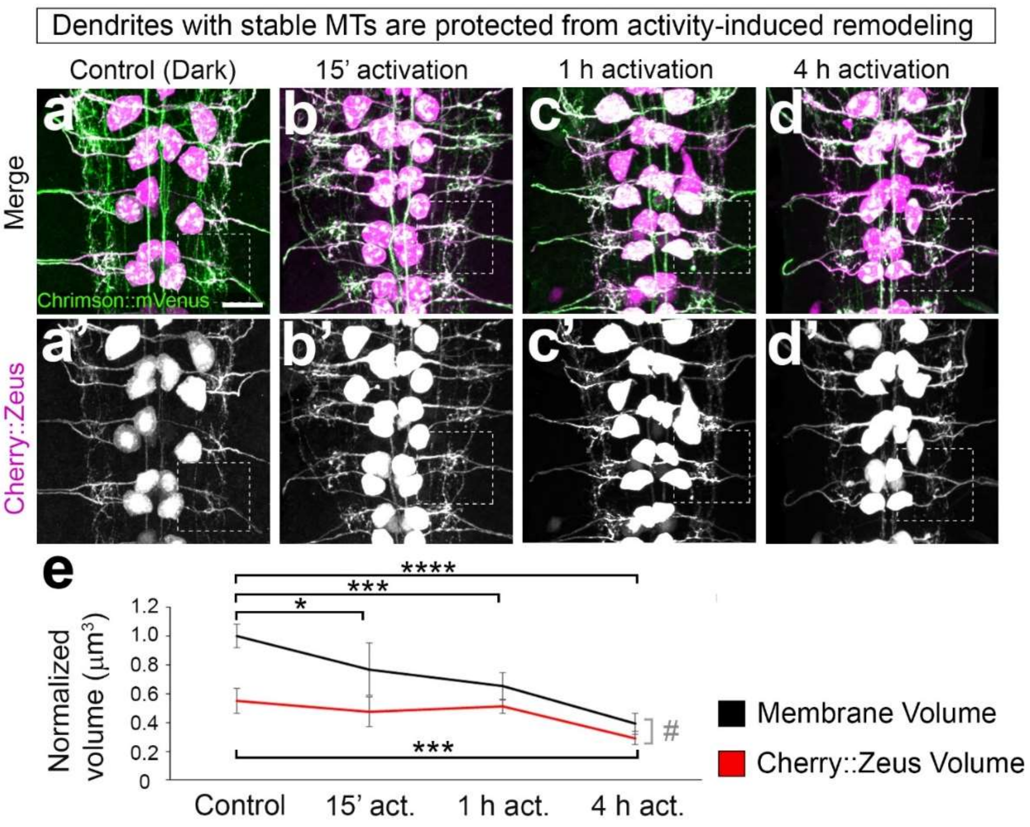


**Extended Data Figure 7. Tubulin stability correlates with dendrite retention during activity-induced remodeling.**

(**a-e**) Dendrites with stable microtubules are resistant to activity-induced remodeling. Representative 3D projections of brains expressing Chrimson (green) and the microtubule reporter Zeus (a tagged microtubule binding protein, magenta) in aCC/RP2 MNs (*RN2-gal4,UAS-Cherry::Zeus,UAS-CsChrimson::mVenus*) at 0 h ALH. Brains were preserved with cold fixative to visualize stable microtubule populations in (**a**) control and after Chrimson-activation for (**b**) 15', (**c**) 1 h, or (**d**) 4 h. Prime panels show Cherry:Zeus channel alone. Scale bar, 10 µm. Boxed in regions represent regions of interest (ROIs) that were used for Imaris "Surface" reconstructions to determine dendrite and microtubule volume.

(**e**) Quantification of the normalized volume of dendrite membranes (Chrimson::mVenus+) and Cherry::Zeus within the same ROI. Microtubule volumes at each time point were calculated relative to the membrane volume for dark-reared controls. The volume per individual brain represents the average volume across 4 hemisegments (A1-A2). In dark-reared controls (N=4), stable microtubule populations reflect 55±8% of the total dendritic volume. Chrimson-activation results in a significant decrease in total dendritic volume after 15' (N=6) and 1 h (N=4) of activation. After 4 h of activation (N=6), both membrane volume and microtubule volume are significantly reduced; however, dendrites with stable microtubules are preferentially retained such that membrane volume is nearly equivalent to the Cherry::Zeus volume (#, $p<.02$, one-way ANOVA).

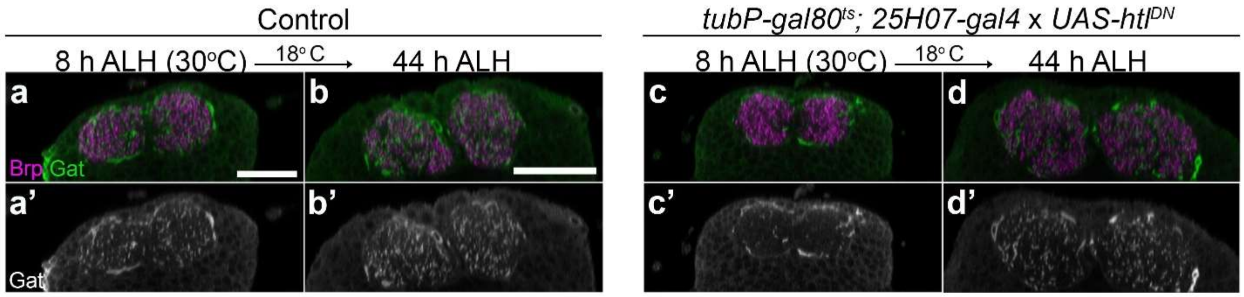


**Extended Data 8. Validation of genetic tool for conditional KD of astrocyte genes.**
(**a-d'**) Orthogonal views through the ventral nerve cord showing the extent of astrocyte infiltration (Gat+, green) into the synapse-dense neuropil (Brp+, magenta). Prime panels show Gat signal alone. (**a-b'**) In control animals (*25H07-gal4* X *UAS-myr::GFP*), astrocytes progressively infiltrate the neuropil from 8 h ALH through 44 h ALH. (**c-c'**) When reared at 30°C through 8 h ALH, expression of *UAS-htl*$^{DN}$ in astrocytes (*tubP-gal80*$^{ts}$*; 25H07-gal4*) suppressed astrocyte infiltration. (**d-d'**) Shifting to 18°C at 8 h ALH resulted in inhibition of Gal4 by TubP-Gal80$^{ts}$, reduced expression of *htl*$^{DN}$, and rescued astrocyte infiltration at 44 h ALH (25°C standard, see methods for details on staging). 8 h scale bar, 20 μm. 44 h scale bar, 30 μm.

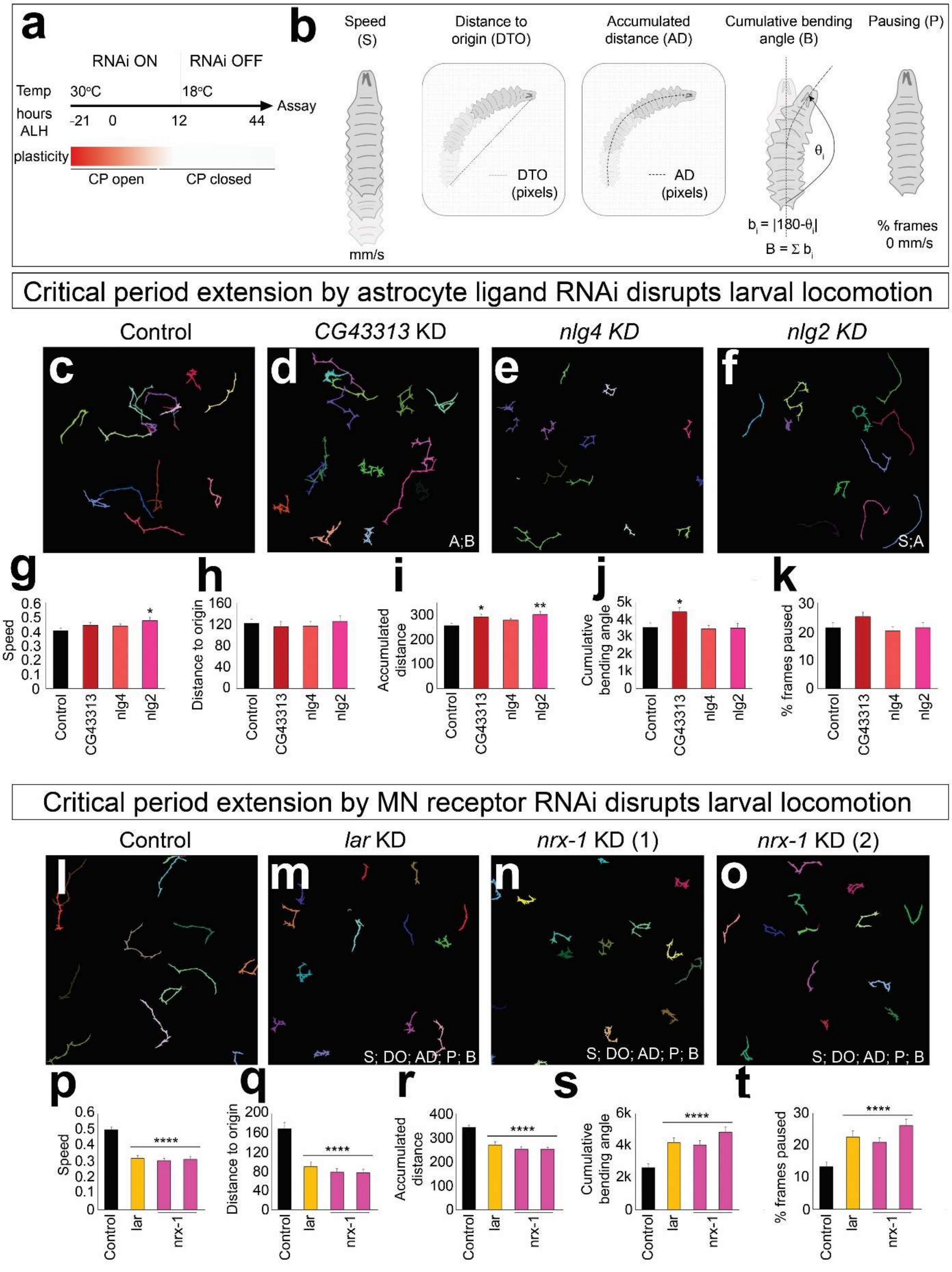
a
RNAi ON
RNAi OFF
Temp
30°C
18°C
hours ALH
-21
0
12
44
Assay
plasticity
CP open
CP closed
b
Speed (S)
mm/s
Distance to origin (DTO)
DTO (pixels)
Accumulated distance (AD)
AD (pixels)
Cumulative bending angle (B)
θi
bi = |180-θi|
B = Σ bi
Pausing (P)
% frames 0 mm/s
Critical period extension by astrocyte ligand RNAi disrupts larval locomotion
Control
CG43313 KD
nlg4 KD
nlg2 KD
c
d
e
f
A;B
S;A
g
h
i
j
k
Speed
Distance to origin
Accumulated distance
Cumulative bending angle
% frames paused
Control
CG43313
nlg4
nlg2
Critical period extension by MN receptor RNAi disrupts larval locomotion
Control
lar KD
nrx-1 KD (1)
nrx-1 KD (2)
l
m
n
o
S; DO; AD; P; B
p
q
r
s
t
Speed
Distance to origin
Accumulated distance
Cumulative bending angle
% frames paused
****
Control
lar
nrx-1

**Extended Data Figure 9. Timely closure of the critical period is required for normal locomotor behavior.**

(**a, b**) Schematic of the experimental paradigm. (**a**) Animals were reared at 30°C from embryo through 12 ± 2 h ALH to extend critical period plasticity, then shifted to 18°C until 44 h ALH (25° staging standard, see Methods). Genotypes: *TubP-gal80$^{ts}$ 25h07-gal4* x *UAS-RNAi* or background control (for astrocyte manipulation); *RN2-gal4; CQ2-gal4* x UAS-RNAi*; TubP-gal80$^{ts}$* for MN manipulation. (**b**) Five behavioral metrics assayed.

(**c-f**) FIMTrack traces from individual freely behaving larvae from (**c**) control and conditional astrocyte-KD of (**d**) *CG43313*, (**e**) *nlg4*, or (**f**) *nlg2*. Traces derived from a one-minute video imaged at 4 Hz. Letters denote behavioral metrics that differed significantly from control.

(**g-k**) Quantification of each behavioral metric for conditional astrocyte KD conditions relative to control. N≥51 larvae per condition. Note that *gat* KD traces are not provided as early KD of *gat* resulted in unrecoverable paralysis.

(**l-o**) FIMTrack traces from individual freely behaving larvae from (**l**) control and conditional MN-KD of (**m**) *lar* or (**n-o**) *nrx-1*. Traces were derived from a one-minute video imaged at 4 Hz.

(**p-t**) Quantification of each behavioral metric in conditional MN-KD conditions relative to control. N≥23 larvae per condition. (**g-k**, **p-t**) Unpaired Student's T-test. Error bars, standard error of the mean.

#### SUPPLEMENTARY MATERIALS: METHODS

##### Lead Contact and Materials Availability

Additional information and inquiries regarding resources and reagents should be directed to and will be fulfilled upon request by Lead Contacts Sarah D Ackerman and Chris Q Doe.

##### Experimental Model and Subject Details

Fly husbandry

All flies were raised at 25°C on standard cornmeal fly food. Animals were staged relative to a 25°C standard. At 25°C, embryos take 21 hours to hatch into larvae[58]; larvae hatch and develop 1.25x faster at 30°C, and 2x slower at 18°C.

Transgenes (in order of appearance)

1. *RN2-gal4* [59]
2. *20XUAS-CsChrimson::mCherry* [60]
3. *20XUAS-CsChrimson::mVenus* (BDSC# 55136)
4. *hs-FLPG5;; 10xUAS(FRT.stop)myr::smGdP-HA, 10xUAS(FRT.stop)myr::smGdP-V5, 10xUAS-(FRT.stop)myr::smGdP-FLAG* (hs-MCFO, BDSC# 64085)
5. *10XUAS-myr::GFP* (BDSC# 32198)
6. *UAS-GtACR2::EYFP* [61]
7. *UAS-shibire*$^{ts1}$ (BDSC# 44222)
8. *UAS-cherry::zeus* [62]
9. *R78F07-lexA* [63]
10. *8xlexAop-2xbrp-short::cherry* [64]
11. *R94E10-lexA* [60]
12. *alrm-lexA::GAD* (Chromosome 3) [65]
13. *lexAop-rpr* [66]
14. *R25H07-gal4* (BDSC# 49145)
15. *13XlexAop2-CsChrimson::mVenus* (BDSC# 55137)
16. *RN2-lexA* (courtesy of Ellie Heckscher, UChicago)
17. *alrm-gal4* (Chromosome 3) [67]
18. *UAS-SAM.dCas9.GS05146* (BDSC# 82741)
19. *UAS-SAM.dCas9.GS04054* (BDSC# 81420)
20. *13lexAop-CsChrimson::tdTomato* (courtesy of Vivek Jayaraman, Janelia Research Campus)
21. *CQ2-gal4* [59]
22. *tubPgal80*$^{ts}$ (BDSC# 7019)
23. *UAS-htl*$^{DN}$ (BDSC# 5366)

Complete list of RNAi lines and other lines used for screening in astrocytes

*gat* RNAi (BDSC# 29422), *CG43313* RNAi (BDSC# 53990, NIG# HMJ-22218), *shot* RNAi (BDSC#s 28366, 64041), *dally* RNAi (BDSC#s 28747, 33952), *nlg4* RNAi (BDSC# 58119, NIG# HMJ-22056), *nlg2* RNAi (BDSC#s 58128, 28331), *kek2* RNAi (BDSC# 31874, NIG# HMJ-21692), *eph* RNAi (BDSC# 28511), *ptp99a* RNAi (BDSC# 25840), *arr* RNAi (BDSC# 53342), *nrx-1* RNAi (BDSC# 27502), *dpy* RNAi (BDSC# 36691), *ft* RNAi (BDSC# 29566), *sparc* RNAi (BDSC# 40885), *mew* RNAi (BDSC# 27543), *sema1b* RNAi (BDSC# 28588), *tsp* RNAi (BDSC# 44116), *CG17739* RNAi (BDSC# 28770), *gbb* RNAi (BDSC# 34898), *rangap* RNAi (BDSC# 29565), *daw* RNAi (BDSC# 34974), *lrch* RNAi (BDSC# 31871), *mav* RNAi (BDSC# 36809), *twin* RNAi (BDSC# 32490), *stat92e* RNAi (BDSC#s 35600, 33637), *eaat1* RNAi (BDSC# 43287), *nlg3* RNAi (BDSC# 38264), *myo* RNAi (BDSC# 36840), *drpr* RNAi (BDSC# 36732), *dpp* RNAi (BDSC#s 33618, 25782), *egr* RNAi (BDSC#55276), *CG16868* RNAi (BDSC# 29617), *tkv* RNAi (BDSC# 35653), *fasII* RNAi (BDSC# 34084), *inx2* RNAi (BDSC# 42645), *lar* RNAi (BDSC# 40938), *7b2* RNAi (BDSC# 27989), *octalpha2* RNAi (BDSC# 50678), *lpr1* RNAi (BDSC# 50737), *egfr* RNAi (BDSC# 36773), *otk* RNAi (BDSC# 25790), *tsp96f* RNAi (BDSC# 40901), *galectin* RNAi (BDSC# 34880), *sfl* RNAi (BDSC#s 34601, 33606), *sgl* RNAi (BDSC# 65348), *trol* RNAi (BDSC#s 29440, 42783, 38298), *tenA* RNAi and overexpression (BDSC#s 42018, 41564), *hSod1* misexpression (BDSC#s 33606, 33607). BDSC: Bloomington Drosophila Stock Center, USA. NIG: Shigen National Institute of Genetics, Japan.

RNAi lines used for screening in aCC/RP2

*lar* RNAi (BDSC#s 40938, 34965), *nrx-1* RNAi (BDSC#s 27502, 32408), *kek2* RNAi (BDSC# 31874), *kek6* RNAi (BDSC# 61212), *kek1* RNAi (BDSC# 57000), *dg* RNAi (BDSC# 34895), *egfr* RNAi (BDSC#s 36773, 25781), *tkv* RNAi (BDSC#s 35166, 33653), *dpp* RNAi (BDSC#s 33618, 25782), *wit* RNAi (BDSC# 25949), *brp* RNAi (BDSC# 25891).

Animal collections

*For live imaging of wildtype samples:* Crosses were reared at 25°C in collection bottles fitted with 3.0% agar apple juice caps containing plain yeast paste. Embryos were then collected on 3.0% agar apple juice caps with plain yeast paste for 1.5 hours and aged at 25°C until hatching.

*For optogenetics* (without RNAi or overexpression (OE))*:* Crosses were reared at 25°C in collection bottles fitted with 3.0% agar apple juice caps containing yeast paste that was supplemented with 0.5mM all-*trans* retinal (+ATR) (Sigma-Aldrich, R2500-100MG). Crosses were supplied fresh yeast paste (+ATR) for a minimum of 72 hours prior to embryo collection to ensure maternal transfer of ATR into embryos. Embryos were then collected on 3.0% agar apple juice caps with yeast paste (+ATR) for 1.5 hours and aged at 25°C. To prevent premature optogenetic activation, crosses and embryos were dark-reared until the appropriate developmental stage. At 25° C, *Drosophila* embryos hatch at 21 h after egg laying (AEL). For assessment of remodeling at 0 h ALH, embryos were aged to 17 h or 20 h AEL prior to light-activation/silencing for 4 h or 1 h, respectively. For 15' manipulations at 0 h after larval hatching (ALH), optogenetic activation/silencing was performed just after hatching. For assessment of activity-induced remodeling at later larval stages, larvae were collected at 0 h ALH and transferred to fresh apple caps (N=20 larvae per cap) containing yeast paste (+ATR), and then manipulated for the designated length of time just preceding the desired larval stage (4, 8, or 22 h ALH). All animals were dissected immediately following activity manipulation.

*For optogenetics* (with RNAi or OE)*:* Crosses were reared at 25°C in collection bottles fitted with 3.0% agar apple juice caps containing yeast paste that was supplemented with 0.5mM all-*trans* retinal (+ATR). Crosses were supplied fresh yeast paste (+ATR) for a minimum of 72 hours prior to embryo collection to ensure maternal transfer of ATR into embryos. Embryos were then collected on 3.0% agar apple juice caps with yeast paste (+ATR) for 1.5 h and aged at 30°C. To prevent premature optogenetic activation, crosses, embryos, and larvae were dark-reared until the appropriate developmental stage. For assessment of activity-induced remodeling post-critical period, larvae were collected at 0 h ALH and transferred to fresh apple caps (N=10 larvae per cap) containing yeast paste (+ATR) and aged 6.5 h (~ 8 h ALH at 25°C standard) prior to dissection for dark-reared controls, or aged 5.5 h followed by 1h light activation. For assessment of activity-induced remodeling with OE, larvae were collected at 0 h ALH and transferred to fresh apple caps (N=10 larvae per cap) containing yeast paste (+ATR) and aged 3.5 h (~ 4 h ALH at 25°C standard) prior to dissection for dark-reared controls, or aged 2.5 h followed by 1 h light activation. All animals were dissected immediately following activity manipulation.

*For thermogenetics via TrpA1*: Crosses were reared at 23°C in collection bottles fitted with 3.0% agar apple juice caps containing plain yeast paste. Embryos were then collected on 3.0% agar apple juice caps with plain yeast paste for 1.5 hours and aged at 23° C. Embryos take 24 h to hatch into larva at 23° C. For assessment of remodeling at 0 h ALH, embryos were aged for 20 h or 23 h AEL prior to thermogenetic activation (4 or 1 h) at 29° C, which induces neuronal firing at ≥ 30 Hz[68].

*For thermogenetics via Shibire$^{ts}$:* Crosses were reared at 25°C in collection bottles fitted with 3.0% agar apple juice caps containing plain yeast paste. Embryos were then collected on 3.0% agar apple juice caps with plain yeast paste for 1.5 hours and aged at 25° C. At 17 h or 19.5 h AEL, Shibire$^{ts}$ embryos were transferred to 30°C to induce silencing for 3h or 1h, respectively. Shibire$^{ts}$ controls were maintained at 25° C. All animals were dissected immediately following activity manipulation.

*For behavioral analyses*: For conditional KD of genes in astrocytes or MNs, we used TubGal80$^{ts}$ as a temperature sensitive repressor of Gal4 (active at 18°C, inactive at 30°C). Crosses were reared at 25°C in collection bottles fitted with 3.0% agar apple juice caps containing plain yeast paste. Embryos were then collected on 3.0% agar apple juice caps with plain yeast paste for 1.5 h and aged at 30°C. Larvae were collected at 0 h ALH and transferred to fresh apple caps (N=40 larvae per cap) supplied with plain yeast and maintained at 30°C for 10 h (~12 ± 2 h ALH at 25°C standard) for robust, Gal4-dependent RNAi expression. Animals were then shifted to 18°C for 64 hours to suppress Gal4 activity and assayed for locomotor defects at ~44 h ALH (25°C standard).

*For astrocyte ablation*: Crosses were reared (*alrm-lexA lexAop-rpr*, astrocyte ablation is Rpr-dependent) at 25°C in collection bottles fitted with 3.0% agar apple juice caps containing yeast paste that was supplemented with 0.5mM all-*trans* retinal (+ATR). Crosses were supplied fresh yeast paste (+ATR) for a minimum of 72 hours prior to embryo collection to ensure maternal transfer of ATR into embryos. To prevent premature optogenetic activation, crosses, embryos, and larvae were dark-reared until the appropriate developmental stage. Larvae were collected at 0 h ALH and transferred to fresh apple caps (N=10 larvae per cap) containing yeast paste (+ATR), and then manipulated for the designated length of time just preceding the desired larval stage (8 h ALH). All animals were dissected immediately following activity manipulation.

#### METHOD DETAILS

##### Optogenetic activation/silencing

The following strategy was used for optogenetic activation/silencing in all fixed preparation experiments. Both Chrimson[69] and GtACR2[61] are activated by a broad spectrum of wavelengths. At the designated stage, dark-reared embryos/larvae (reared on apple caps with yeast + ATR) were placed beneath a full spectrum light bulb, shining with an average intensity of 10550 lx (determined with two independent photometer software programs developed for Android: Light Meter© and Photometer©). Animals were dissected in low-light conditions (<100 lx) immediately following activity manipulations. For recovery experiments, animals were transferred back into dark-rearing conditions at 25°C until dissected in low-light conditions. We used tonic activating conditions, which occur *in vivo* in a number of mutant models[58,70], to drive the homeostatic response.

##### MultiColor FlpOut Clone Generation

At 15 h AEL, embryos were prepared for heatshock to induce FLP-out clones. Apple caps covered in embryos were sliced to a thickness of ~2 mm and then adhered to a 100 X 26 mm petri dish with water to increase heat transfer. The petri dish was then sealed with parafilm and floated on a 37°C water bath for 17 minutes, followed by 15 minutes at 18°C to prevent further FLP-out events. Embryos were then transferred back to 25°C until the designated stage and/or manipulation.

Immunohistochemistry

Larval brains were dissected in sterile-filtered, ice-cold 1X PBS and mounted on 12mm #1 thickness poly-D-lysine coated round coverslips (Corning® BioCoat™, 354085). Brains were fixed for either 12 minutes (0-8h ALH samples) or 15 minutes (22h ALH samples) in fresh 4% paraformaldehyde (Electron Microscopy Sciences, 15710) in .3% PBSTriton, and then washed in .3X PBSTriton to remove fixative. Samples were blocked overnight at 4°C in .3% PBSTriton supplemented with 1% BSA (Fisher, BP1600-100), 1% normal donkey serum and 1% normal goat serum (Jackson ImmunoResearch Laboratories, Inc., 017-000-121 and 005-000-121). Brains were then incubated in primary antibody for one-two days at 4°C. The primary was removed, and brains were washed overnight at 4°C with 0.3% PBST. Brains were then incubated in secondary antibodies overnight at 4°C. The secondary antibodies were removed, and brains transferred to .3% PBSTriton overnight prior to DPX mounting. Brains were dehydrated with an ethanol series: 30%, 50%, 70%, 90%, each for 5 minutes, then twice in 100% ethanol for 10 minutes each (Decon Labs, Inc., 2716GEA). Finally, samples were incubated in xylenes (Fisher Chemical, X5-1) for 2 x 10 minutes, were mounted onto slides containing DPX mountant (Millipore Sigma, 06552), and cured for 1-2 days before imaging.

The following primary and secondary antibodies were used:

| Primary Antibody (concentration) | Source | Figures |
|---|---|---|
| Mouse anti-Cherry (1:500) | Clontech Cat. 632543 | 1- 4; ED 1,5 |
| Chicken anti-GFP (1:1000) | Aves Cat. GFP-1010 | 1,4; ED 1,7 |
| Guinea Pig anti-GFP (1:500) | Frontier Institute Cat. GFP-GP-Af1180 | 2; ED 4 |
| Rabbit anti-V5 (1:500) | Cell Signaling Technology Cat. 13202 | 2; ED 2,6 |
| Rat anti-HA (1:100) | Millipore Sigma Cat. 11867423001 | 2; ED 2,6 |
| Rabbit anti-cherry (1:500) | Novus Biologicals, Cat. NBP2-25157 | 2,4; ED 4,7 |
| Rabbit anti-Gat (1:4000) | M. Freeman lab | 3,4; ED 6,8 |

| | | |
|---|---|---|
| Rabbit anti-GABA (1-500) | Sigma A2052 | ED 4 |
| Rabbit anti-dsred (1:500) | Takara Bio Cat. 632496 | 4 |
| Ms anti-Brp/Nc82 (1:100) | DSHB nc82 | ED 8 |

Secondary Antibodies:
All secondary antibodies were purchased from Jackson ImmunoResearch and used at a working concentration of 1:400. The following antibodies were used: Alexa Fluor® Rhodamine RedTM-X Donkey-Anti Mouse (715-295-151), Alexa Fluor® 488 Donkey anti-Guinea Pig (706-545-148), Alexa Fluor® 488 Donkey anti-Chicken (703-545-155), Alexa Fluor® Rhodamine RedTM-X Donkey-Anti Rat (712-295-153), Alexa Fluor® 488 Donkey-Anti Rat (712-545-153), Alexa Fluor® 647 Donkey-Anti Rb (711-605-152); Alexa Fluor® 488 Donkey anti-Rabbit (711-545-152), Alexa Fluor® Rhodamine RedTM-X Donkey Anti-Rabbit (711-295-152).

Light Microscopy
Fixed larval preparations for dendrite and astrocyte morphology analyses were imaged with a Zeiss LSM 700 laser scanning confocal using a 63x/1.4 NA Oil Plan-Apochromat DIC m27 objective lens. Fixed larval preparations for synapse quantifications were imaged on a Zeiss LSM 800 laser scanning confocal fitted with a 63x/1.40 NA Oil Plan-Apochromat DIC m27 objective lens and GaAsP photomultiplier tubes.

Image processing and analyses
Quantitative analyses (Figures 1-4, Extended Data Figures 2-4,6-7) were performed using Imaris 9.2.0 (Bitplane AG). Visualization and projection of images for phenotypic categorization (Figures1,3, Extended Data Figure 1,5) were performed in FIJI (ImageJ 1.50d, https://imagej.net/Fiji).

Time-lapse imaging of fictive preparations
The following assay was used for live imaging of aCC/RP2 dendrites in isolated CNS (Figures 3-4, Extended Data Figure 3). Larvae were dissected at the indicated stage in a hemolymph-like solution (HL3.1); both lobes and the ventral nerve cord were kept intact. Isolated brains were placed on a 12mm #1 thickness poly-D-lysine coated round coverslips (Corning® BioCoat™, 354085), a single 18mm x 18mm x 0.16mm cover glass (Fisher, 12-542B) was pressed against the coverslip until brain lobes were slightly compressed, a drop of HL3.1was used to facilitate sealing. For optogenetic experiments, both dissections and mounting were performed under low light conditions (<100 lx) to delay optogenetic activation for Chrimson experiments and controls. For imaging, we utilized Zeiss LSM 800 laser scanning confocal fitted with a 63x/1.40 NA Oil Plan-Apochromat DIC m27 objective lens and GaAsP photomultiplier tubes. For Figure 3 and Extended Data Figure 3, continuous

scans were obtained every 45 seconds, for 15 minutes, with two hemisegments in the field of view. A z-stack of 25 μm (allowing for drift in Z) with 1 μm step size was performed for Chrimson experiments, and .5 μm for time course analyses. For microtubule dynamics during dendrite retraction (Figure 4), a z-stack of 21 μm (allowing for drift in Z) with 0.3 μm step size was imaged continuously with stacks acquired every 10 seconds for 10 minutes, with only one dendritic branch in the field of view.

Behavioral analyses

Larvae were transferred from 3.0% agar caps with yeast to 1.2% agarose plates for half an hour prior to locomotion assays to avoid inadvertent, temperature-dependent changes in locomotion and to allow the larvae to acclimate to the new crawling surface. Larvae were then transferred to a FIM behavior table[71] fitted with a fresh 1.2% agarose gel and allowed to further acclimate for two minutes prior to imaging. Two or more independent cohorts of larvae (N=15 larvae per cohort) were tested per genotype. Larval crawling was imaged at 4 Hz, 91 pixels/cm for one minute using a Basler acA2040-25gm camera in the Pylon5 Camera Software Suite (Basler). Data were then analyzed using FIMTrack software using standard settings[71].

Figure preparation

Images in figures were prepared as either 3D projections in Imaris 9.2.0 (Bitplane AG) or 3D projections in FIJI (ImageJ 1.50d) and assembled using Adobe Illustrator or Adobe Photoshop. Schematics were drawn in Microsoft Powerpoint.

#### QUANTIFICATION AND STATISTIC ANALYSIS

*Phenotypic categorization*

For classification of post-activation dendrite morphologies as either control, mildly reduced, or strongly reduced (Figures 1,3, Extended Data Figure 1,5), 3D projections of data were generated in Fiji to standardize sample angle (dorsal up, anterior top) and fluorescence intensity. Projections were then blinded to genotype and phenotype and scored phenotypically.

*Volumetric assays*

For quantification of dendrite volume for critical period analysis (Figures 1,4, Extended Data Figure 7), data was acquired with a voxel size of 0.124 x 0.124 x 0.325 $\mu m^3$. aCC/RP2 dendrites within a single hemisegment (A1-A2) were captured in a standard ROI spanning 100 pixels by 100 pixels in XY and 8.125 μm in Z with the top of the ROI beginning dorsally at the aCC/RP2 axons for 0h, 4h, and 8h ALH analyses. The ROI was adjusted to account for increased brain size at 22h ALH: 100 pixels by 150 pixels in XY and 8.125 μm in Z at 22h ALH. Dendrites were then reconstructed using the Imaris "Surface" module using default thresholding to determine the total dendritic volume within the ROI. Dendrite volume per brain was determined by averaging the dendrite volume of 4 individual hemisegments. Chrimson data was normalized to time-matched, experiment-matched, dark-reared controls. For quantification of astrocyte volume (Extended Data 6), data was acquired with a voxel size of 0.397 x 0.397 x 0.427 $\mu m^3$. A ROI was built around each individual astrocyte (varying sizes) and astrocytes were then reconstructed using the Imaris "Surface" module using default thresholding to determine the total astrocyte volume within the ROI.

*Single RP2 reconstructions*
For morphometric analyses of individual RP2 MN MCFO clones (Figure 2, Extended Data Figure 2), data were acquired with a voxel size of .099 x .099 x .318 $\mu m^3$. Single RP2 MNs were captured within a ROI and reconstructed using the Imaris "Filaments" function (starting position: base of cell body; largest diameter filament: 3 µm; seed points: .2 µm; thresholds varied with fluorescence intensity). Reconstructions were adjusted manually when necessary.

*Synapse quantification*
Data were acquired with a voxel size of .076 x .076 x .27 $\mu m^3$ and de-convoluted in Imaris. Chrimson::mVenus+ aCC/RP2 dendrites within a single hemisegment were reconstructed using the Imaris "Surface" module (no smoothing, thresholds varied with fluorescence intensity). A standard ROI spanned 150 x 150 pixels in XY, and 6.75 µm in Z with the top of the ROI beginning dorsally at the aCC/RP2 axons. A distance transformation was then performed on the "Surface". Brp-Short::Cherry+ presynapses within the ROI were annotated using the Imaris "Spots" functions and then classified as "direct" synapses if they fell within 90 nm of the "Surface" based on previously validated criteria[72].

<u>Analysis of time-lapse imaging samples</u>
Images were corrected for 3D drift using a Fiji 3D correction plug-in from the Imaris interface, as well as corrected for bleaching using the 'histogram matching' function in the same software. Data were then imported to Imaris and analyzed using the "Filaments" module. An automatic detection (starting position: base of cell body; largest diameter filament: 2 µm; seed points: ~.15 µm; thresholds varied with fluorescence intensity) was used for the first time point. Reconstruction of subsequent time points was done manually, using the initial automatic reconstruction as a guide to ensure correct 3D position across timepoints ("cone" reconstruction was set to .3 µm). For quantification of dendrite length extension/retraction events, only dendrites present throughout the 15 minutes were analyzed. For Chrimson experiments, post-activation dendrite lengths were normalized to pre-activation dendrite length to assess retraction over time. Lengths were then compartmentalized to 10 normally distributed values using MATLAB (Mathworks) to minimize variations in brain size and process length between WT and Chrimson-induced activation. Both populations were then plotted against each other as a function of time. For

dendrite dynamicity studies, average displacement quantification was calculated per brain (N=5 per timepoint), encompassing 10 reconstructed dendritic processes each. An extension/retraction event was given a value of 1, stability at each time frame was given a value of 0. Motility of a dendrite was defined as an “event” when a length difference of 0.50 μm was detected compared to the previous timepoint. “Motile dendrite” was defined as at least one filopodial event over the 15-min experiment. For quantification of microtubule collapse, retracting dendritic filopodia were identified using Imaris 3D viewer. A retraction event was defined as a ≥0.75 μm change in filopodial length within a period of 40 seconds. The lower boundary of the ROI was created in the retracting process using the final filopodial length as a landmark at t=40 seconds. The upper boundary of the ROI spanned from the landmark to the most distal length of the filopodial process at $T_0$. Channel intensity was calculated using the “intensity sum” value within Imaris's “surface” function.

Statistical analyses

Statistics were performed using a combination of Microsoft Excel, MATLAB (MathWorks), and Prism (GraphPad) software. One-way ANOVA was used unless otherwise noted. Error bars, Standard Deviation unless otherwise noted. A 95% confidence interval was used to define the level of significance. Significance: *, p<.05; **, p<.01; ***, p<.001; ****, p<.0001, NS= not significant. ♦ used in place of * to denote significance following two-way ANOVA when both one-way and two-way ANOVA data are displayed on a single graph. All other pertinent information, including sample size, statistical test employed, and variance can be found in the figure legends or labeled within the figure.

Methods References

58 Crisp, S., Evers, J. F., Fiala, A. & Bate, M. The development of motor coordination in Drosophila embryos. *Development (Cambridge, England)* **135**, 3707-3717, doi:10.1242/dev.026773 (2008).
59 Landgraf, M., Jeffrey, V., Fujioka, M., Jaynes, J. B. & Bate, M. Embryonic origins of a motor system: motor dendrites form a myotopic map in Drosophila. *PLoS Biol* **1**, E41, doi:10.1371/journal.pbio.0000041 (2003).
60 Carreira-Rosario, A. *et al.* MDN brain descending neurons coordinately activate backward and inhibit forward locomotion. *eLife* **7**, doi:10.7554/eLife.38554 (2018).
61 Mohammad, F. *et al.* Optogenetic inhibition of behavior with anion channelrhodopsins. *Nat Methods* **14**, 271-274, doi:10.1038/nmeth.4148 (2017).
62 Volpi, S., Bongiorni, S., Fabbretti, F., Wakimoto, B. T. & Prantera, G. Drosophila rae1 is required for male meiosis and spermatogenesis. *Journal of cell science* **126**, 3541-3551, doi:10.1242/jcs.111328 (2013).
63 Zarin, A. A., Mark, B., Cardona, A., Litwin-Kumar, A. & Doe, C. Q. A multilayer circuit architecture for the generation of distinct locomotor behaviors in Drosophila. *eLife* **8**, doi:10.7554/eLife.51781 (2019).

64 Berger-Muller, S. *et al.* Assessing the role of cell-surface molecules in central synaptogenesis in the Drosophila visual system. *PloS one* **8**, e83732, doi:10.1371/journal.pone.0083732 (2013).
65 Stork, T., Sheehan, A., Tasdemir-Yilmaz, O. E. & Freeman, M. R. Neuron-glia interactions through the Heartless FGF receptor signaling pathway mediate morphogenesis of Drosophila astrocytes. *Neuron* **83**, 388-403, doi:10.1016/j.neuron.2014.06.026 (2014).
66 Herranz, H., Weng, R. & Cohen, S. M. Crosstalk between epithelial and mesenchymal tissues in tumorigenesis and imaginal disc development. *Curr Biol* **24**, 1476-1484, doi:10.1016/j.cub.2014.05.043 (2014).
67 Doherty, J., Logan, M. A., Tasdemir, O. E. & Freeman, M. R. Ensheathing glia function as phagocytes in the adult Drosophila brain. *The Journal of neuroscience : the official journal of the Society for Neuroscience* **29**, 4768-4781, doi:10.1523/jneurosci.5951-08.2009 (2009).
68 Pulver, S. R., Pashkovski, S. L., Hornstein, N. J., Garrity, P. A. & Griffith, L. C. Temporal dynamics of neuronal activation by Channelrhodopsin-2 and TRPA1 determine behavioral output in Drosophila larvae. *J Neurophysiol* **101**, 3075-3088, doi:10.1152/jn.00071.2009 (2009).
69 Klapoetke, N. C. *et al.* Independent optical excitation of distinct neural populations. *Nat Methods* **11**, 338-346, doi:10.1038/nmeth.2836 (2014).
70 Peng, J. J. *et al.* A circuit-dependent ROS feedback loop mediates glutamate excitotoxicity to sculpt the Drosophila motor system. *eLife* **8**, doi:10.7554/eLife.47372 (2019).
71 Risse, B. *et al.* FIM, a novel FTIR-based imaging method for high throughput locomotion analysis. *PloS one* **8**, e53963, doi:10.1371/journal.pone.0053963 (2013).
72 Sales, E. C., Heckman, E. L., Warren, T. L. & Doe, C. Q. Regulation of subcellular dendritic synapse specificity by axon guidance cues. *eLife* **8**, doi:10.7554/eLife.43478 (2019).
